# Mapping yeast mitotic 5’ resection at base resolution reveals the sequence and positional dependence of nucleases *in vivo*

**DOI:** 10.1101/2021.02.27.433206

**Authors:** Dominic Bazzano, Stephanie Lomonaco, Thomas E. Wilson

**Affiliations:** Department of Pathology, University of Michigan, Ann Arbor, 48109; Department of Human Genetics, University of Michigan, Ann Arbor, 48109

**Keywords:** DNA repair, double strand break, resection, recombination, mitotic

## Abstract

Resection of the 5’-terminated strand at DNA double strand breaks (DSBs) is the critical regulated step in the transition to homologous recombination. Biochemical and genetic studies have led to a multi-step model of DSB resection in which endonucleolytic cleavage mediated by Mre11 in partnership with Sae2 is coupled with exonucleolytic cleavage mediated by redundant pathways catalyzed by Exo1 and Sgs1/Dna2. These models have not been well tested at mitotic DSBs *in vivo* because most methods used to monitor resection cannot precisely map early cleavage events. Here we report resection monitoring with high-throughput sequencing in which molecular identifiers allow exact counting of cleaved 5’ ends at base pair resolution. Mutant strains, including *exo1*Δ, *mre11*-H125N, *exo1*Δ and *exo1*Δ *sgs1*Δ, revealed a major Mre11-dependent cleavage position 60 to 70 bp from the DSB end whose exact position depended on local sequence and tracked a possible motif. They further revealed an Exo1-dependent pause point approximately 200 bp from the DSB. Suppressing resection extension in *exo1*Δ *sgs1*Δ yeast exposed a footprint of regions where cleavage was restricted within 119 bp of the DSB. These results provide detailed *in vivo* support of prevailing models of DSB resection and extend them to show the combined influence of sequence specificity and access restrictions on Mre11 and Exo1 nucleases.

## Introduction

Accurate and efficient repair of DNA double-strand breaks (DSBs) is critical to cell survival and proper genomic function (Scully et al., 2019). Two evolutionarily conserved pathways exist to repair DSBs: non-homologous end joining (NHEJ) and homologous recombination (HR) (Chapman et al., 2012; Zhao et al., 2020). In NHEJ, DSB termini are rapidly bound by the Ku heterodimer, a ring-like protein complex comprised of Yku70 and Yku80 in yeast (Emerson and Bertuch, 2016). Ku binding constrains nuclease activity at DSB ends and promotes the association of downstream proteins critical to repair (Frit et al., 2019). The Mre11-Rad50-Xrs2 (MRX) complex also appears at DSBs nearly instantaneously and further promotes downstream signaling, such as Tel1-mediated checkpoint activation (Cassani et al., 2019; Myler et al., 2017). MRX also has functions in tethering the two DSB termini and end processing during NHEJ (Käshammer et al., 2019; Williams et al., 2007). In NHEJ, the two DSB ends are ultimately ligated by DNA ligase IV (Dnl4 in yeast) (Wilson et al., 1997; Zhang et al., 2007).

Unlike NHEJ, HR relies on nucleolytic processing of DSB 5’-terminated strands in a mechanism called end resection. This highly regulated process creates an exposed 3’-terminated strand suitable for donor pairing and extension (Symington, 2016). End resection is proposed to begin through a targeted endonucleolytic incision by Mre11 <100 bp away from the DSB end in a reaction that is dependent on Rad50 and Sae2 in yeast (Cannavo and Cejka, 2014; Käshammer et al., 2019). After this incision, Mre11 uses its 3’-5’ exonucleolytic activity to degrade DNA back towards the DSB end, destabilizing Ku (Chanut et al., 2016). Long range resection in the 5’-3’ direction occurs through Mre11-mediated recruitment of the exonuclease Exo1 or the helicase Sgs1 in partnership with the endonuclease Dna2 (Cejka, 2015; Mimitou and Symington, 2008). The combined activities of Mre11 backtracking toward the DSB end and Exo1-or Sgs1-Dna2-mediated degradation of DNA away from the break comprises the prevailing bidirectional model of resection (Garcia et al., 2011). The resulting ssDNA is ultimately coated by Rad51 to promote HR (Bonilla et al., 2020).

While many proteins involved in HR and resection have been identified, there is still much to be learned about the precise coordination of the earliest initiating events as they occur *in vivo*. Prior *in vitro* studies on which current models are based analyzed Mre11 nuclease activity on Ku-occluded synthetic dsDNA ends and characterized a primary incision ∼35 bp from the terminus on a 70 bp DNA substrate (Reginato et al., 2017). Wang *et al*. also observed a second cleavage event occurring further downstream at 65 bp from the DNA end in addition to the 35 bp incision when the DNA substrate length was increased to 100 bp (Wang et al., 2017). Such work led to a hypothesis that Mre11 makes stepwise endonucleolytic incisions from DSB termini to initiate resection (Cannavo et al., 2019).

Evidence directly supporting such highly precise models of resection is lacking for mitotic DSBs *in vivo*. Most current methods used to monitor resection in *in vivo* typically use qPCR-or Southern Blot-based approaches to quantify ssDNA appearance beginning 100s of bp from the DSB end (Peng et al., 2021; Zhou et al., 2014), outside of the window examined by *in vitro* studies. These methods are highly informative regarding resection kinetics but lack the resolution needed to understand the exact sequence of events during resection initiation, specifically regarding Mre11 and Exo1 nuclease activity and the factors that promote or constrain their patterns of action, including the precise locations of Mre11 incision points. The covalent linkage of Spo11 to meiotic DSB 5’ termini has permitted more detailed pictures of this process but these advantages are not available for mitotic DSBs (Mimitou and Keeney, 2018; Mimitou et al., 2017).

We developed a novel *in vivo* resection assay to map mitotic end resection at a site-specific DSB in budding yeast at base pair resolution. High molecular weight DNA was subjected to primer extension and ligation into circles in a manner that allowed exact counts of the molecules terminated at all possible resection positions within ∼350 bp of the DSB. Loss of Exo1 resulted in increased signal throughout that window, most notably at a position −65/-66 bp from the DSB end that proved to be Mre11-dependent, in explicit support of prevailing resection models (Wang et al., 2017). This pattern was not rigidly positioned but instead moved at different DSB ends in a manner that suggested a possible sequence motif preference, nor was it singular in that less efficient incision points were also observed. A distinct signal peak at a position −206 bp from the DSB was Exo1-dependent and suggested a pause point in progressive resection. The pattern in *exo1 sgs1* mutant yeast exposed a “footprint” of factors restricting cleavage activity near the DSB. Other mutants revealed dynamic activities at the extreme DSB terminus. These studies provide a precise view of the nuclease activities at DSB ends in living mitotic cells and of the sequence-and position-dependent factors that constrain them.

## Results

### Mapping end resection intermediates in NHEJ-deficient yeast

The DSB system was previously described (Wu et al., 2008) and includes (i) the HO endonuclease coding sequence placed under control of the native *GAL1* promoter and (ii) a single HO cut site (HOcs) in a nucleosome-free region of the *ILV1* promoter (Supplementary Figure 1A). To facilitate resection monitoring without competing NHEJ, we added a *dnl4*-K466A mutation to all strains that renders Dnl4 catalytically inactive but able to bind normally to DSBs (Chiruvella et al., 2013), henceforth referred to as wild-type (WT) with respect to DSB resection. These alleles result in rapid and irreversible generation of a site-specific DSB at *IVL1* in >90% of cells within 30 minutes of adding galactose (Figure 1A, Supplementary Figure 2). At 45 minutes after DSB induction, cultures were resuspended in dextrose media to suppress HO expression and stimulate resection (Lomonaco et al., 2021; Wu et al., 2008) as measured in Figure 1B using a ddPCR method based on the resistance of resected ssDNA to digestion by a restriction endonuclease (RE) (Zhou et al., 2014). Care was taken to extract high molecular weight genomic DNA (gDNA) given that sheared gDNA is indistinguishable from resection intermediates (Supplementary Figure 3).

**Figure 1.**
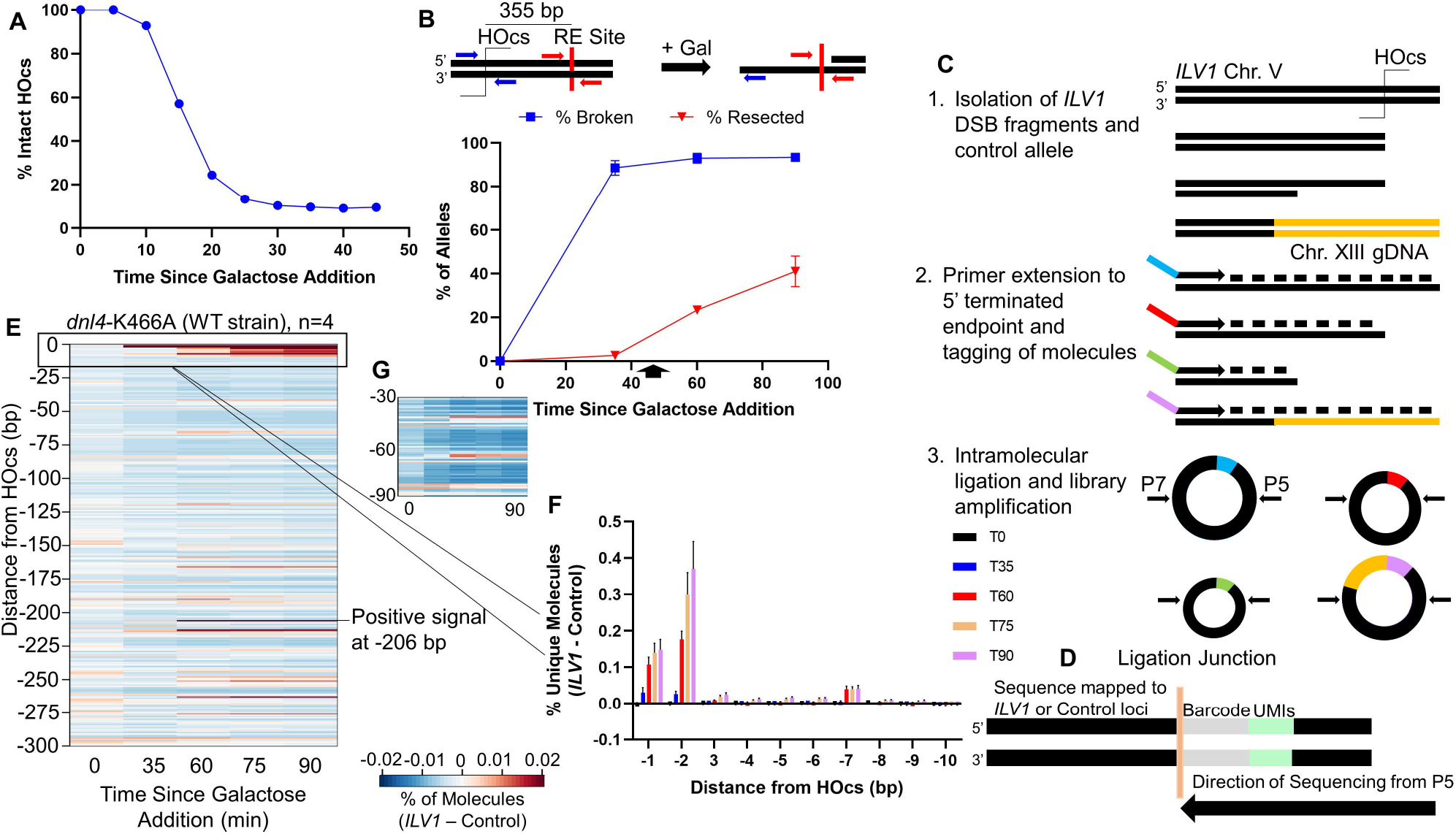
Description and validation of resection sequencing by primer extension and circularization. **(A)** *ILV1* DSB formation efficiency as determined by ddPCR after galactose addition at time 0 to induce HO expression. **(B)** ddPCR-based resection assay to determine the percentage of ssDNA after DSB induction at a site 355 bp away from the HOcs. Cells were washed into glucose at 45 minutes (arrow). **(C)** Steps in resection sequencing to map 5’ terminated end intermediates at base pair resolution using primer extension to add UMIs, followed by intramolecular circularization and PCR of junctions. **(D)** Library products were sequenced such that an indexed P5 primer read, in order, the UMI, a fixed time point barcode and genomic bases from the *ILV1* DSB or a control locus. **(E)** Heatmap depicting the fraction of unique molecules with 5’ endpoints at each position on one side of the *ILV1* HOcs, expressed as a normalized signal above the background fraction of reads aligning to the control locus at similar fragment sizes (see Methods). The NHEJ-defective Dnl4 catalytic point mutant *dnl4*-K466A is considered WT with respect to resection. **(F)** Bar graph showing the percentage of unique *ILV1* molecules with 5’ ends within the first 10 bp away from the HOcs for each time point in WT. Negative position designations indicate the number of bases removed from the 5’ terminated strand. Bars are the average +/-standard deviation of four biological replicates. **(G)** Expansion of the region −30 to −90 bp from the HOcs in WT. The color scale is 2-fold more sensitive than (E).

We further added a control allele to our strains that lacked a DSB but had (i) a shared common sequence with the DSB allele and (ii) a unique sequence abutting an NdeI restriction site, the same site as found 84 bp distal to the *ILV1* HOcs as well as within the shared common sequence (Figure 1C, panel 1, Supplementary Figure 1B,C). Purified gDNA was digested with NdeI followed by annealing and extension of a primer that matched the NdeI site in the shared common sequence, which incorporated a unique molecular identifier (UMI) and a time point barcode into each gDNA molecule (Figure 1C, panel 2, Supplementary Figure 1B-D). Intramolecular ligation created a covalent bond between the UMIs and the 5’ ends of both DSB and control DNA molecules, which might have been created by NdeI, HO, 5’ resection or random shearing (Figure 1C, panel 3). P5 and P7 sequences for high throughput sequencing were added by PCR amplification of pooled ligation products to minimize batch effects (Figure 1D). Because the primer extension primer had a 5’ hydroxyl, only gDNA strands with 5’ phosphates could give final products.

After sequencing, unique source molecules were computationally assigned to the mappable positions of the DSB and control alleles (see Methods). An average of 21M (range 8M to 34M) unique source molecules were obtained per time point (Supplementary File 1). We could not reliably quantify molecules matching the DSB HO and NdeI cleavage positions due to UMI saturation but could quantify the control locus because we used a mixture in which only 5% of cells carried the control allele. Both loci had a minimal background signal at internal positions before the DSB was formed (Supplementary Figure 4). At 35 minutes, signal at the DSB allele shifted to the HOcs position, referred to as position 0 (not shown), and at further time points we observed resection signal above background at positions internal to only the DSB allele (Supplementary Figure 4), referred to with negative numbers to indicate how many bases had been removed from the HO cut end. For visualization, we subtracted the control allele background signal from the DSB allele signal and constructed heat maps (Figure 1E).

### Signal accumulations are evident at distinct positions even in wild-type yeast

Even though resection initiation through our ∼350 bp measurement window is expected to be efficient in our WT strain (Figure 1B), we observed intriguing signal patterns that are explored below. Molecules corresponding to the first few base positions near the HOcs progressively increased from 60 to 90 minutes after DSB induction (Figure 1E-F). Since the DSB had formed by 35 minutes, this increasing signal at 60 minutes must be due to end processing associated with attempted NHEJ or resection initiation. There was also a specific accumulation of signal only in the resecting time points at a single −206 position (Figure 1E). Finally, when we looked near the −30 and −60 positions suggested as Mre11 cleavage positions by the biochemical experiments of Cejka and Sung, we noted a slight increase in signal at selected positions, especially at −65/-66 (Figure 1E). This signal was weak but compelling due to its location, similar time course as above, and the negative footprint surrounding it where very little cleavage was observed (Figure 1G).

### Exo1 loss shifts the DSB end resection pattern more than the loss of Sgs1

To add information to the signal patterns above we repeated resection sequencing in strains lacking Exo1 or Sgs1, which are required for the two main long-range resection mechanisms (Mimitou and Symington, 2008; Zhu et al., 2008). While the *sgs1*Δ resection profile looked largely like that of WT (Figure 2A), *exo1*Δ first led to a decrease in reads aligning near the HOcs, especially at the −2 position, consistent with Exo1 itself being responsible for removing a limited number of terminal bases (Figure 2A-B) (Gobbini et al., 2018). In marked contrast, we observed a dramatic increase in reads mapping to the −65/-66 bp position in the absence of Exo1 (Figure 2A,C), indicating that Exo1 did not create this signal but instead was responsible for suppressing it in WT cells. There was minimal signal in *exo1*Δ after the −65/-66 bp position until −119 bp from the DSB, after which we observed a steady increase in intermediates throughout the sequencing window. Despite this general increase in reads mapping further away from the DSB terminus in *exo1*Δ, loss of Exo1 led to a specific reduction in read accumulation at the −206 bp position, identifying it as a likely pause point for progressive Exo1-mediated resection (Figure 2A,D). The large difference when comparing *sgs1*Δ and *exo1*Δ resection patterns suggests that Exo1-mediated resection is the preferred mode of end resection initiation near the DSB.

**Figure 2.**
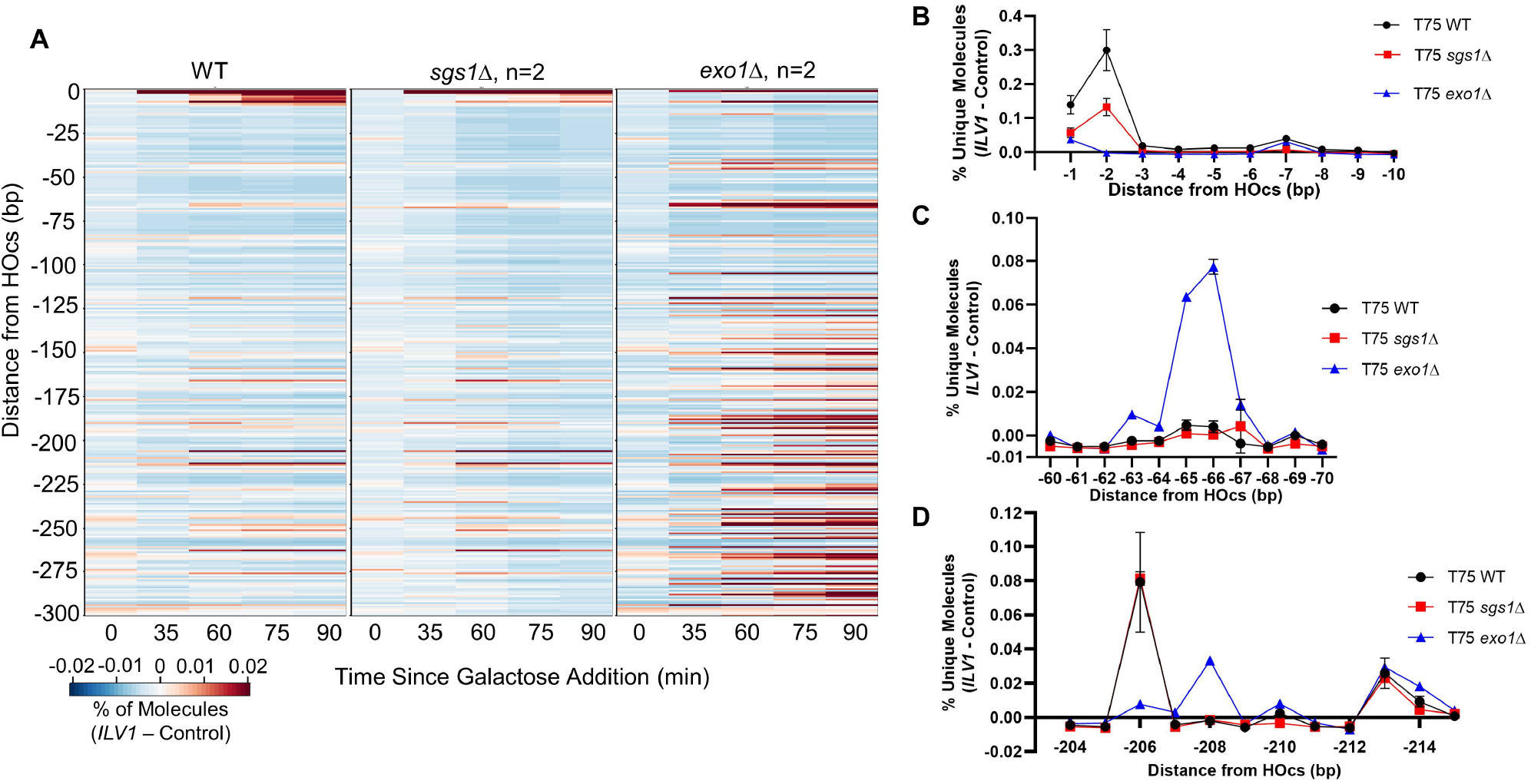
Loss of Exo1 increases resection signal close to the *ILV1* DSB, especially at −65/-66 bp. **(A)** Heatmaps of resection signal similar to Figure 1E for WT, *exo1*Δ and *sgs1*Δ yeast. **(B-D)** The normalized fractional resection signal above background is plotted for WT, *sgs1*Δ and *exo1*Δ yeast at positions **(B)** −1 to −10 bp, **(C)** −60 to −70 bp and **(D)** −204 to −215 bp from the *ILV1* HOcs on the promoter side of the gene. Exo1 loss had opposite effects on the signal obtained at the −2 and −206 positions as compared to the −65/-66 bp peak.

### Mre11 is responsible for the peaks at the −65/-66 and other positions

Results above demonstrate that a protein other than Exo1 must create the signal peak at the −65/-66 position, which prevailing models predict should be Mre11. To explore this, we first examined strains with a complete deletion of *MRE11*. Sensitive examination surrounding the −65/-66 position similar to Figure 1G revealed that loss of Mre11 erased the small signal peak observed in WT cells, although intriguingly the negative footprint surrounding that position remained even without Mre11 present (Figure 3A). With the benefit of Mre11 loss we could also discern that less intense signal peaks were present in WT at −42, −84, −105 and −119 (Figure 3A). Here, it is important to note that our method can only map a single 5’ terminal position of a source molecule relative to the DSB-distal primer. Thus, we cannot definitively judge how effectively the more DSB-proximal −42 position is cleaved relative to the efficient −65/-66 position, since cleavage at −65/-66 would prevent detection of simultaneous cleavage at −42. We can confidently state that the −65/-66 position is cleaved more efficiently than the more DSB-distal positions. Importantly, *mre11*Δ also reduced most signal in our interrogation window after the first ∼10 bp, including the −206 peak (Figure 3B).

**Figure 3.**
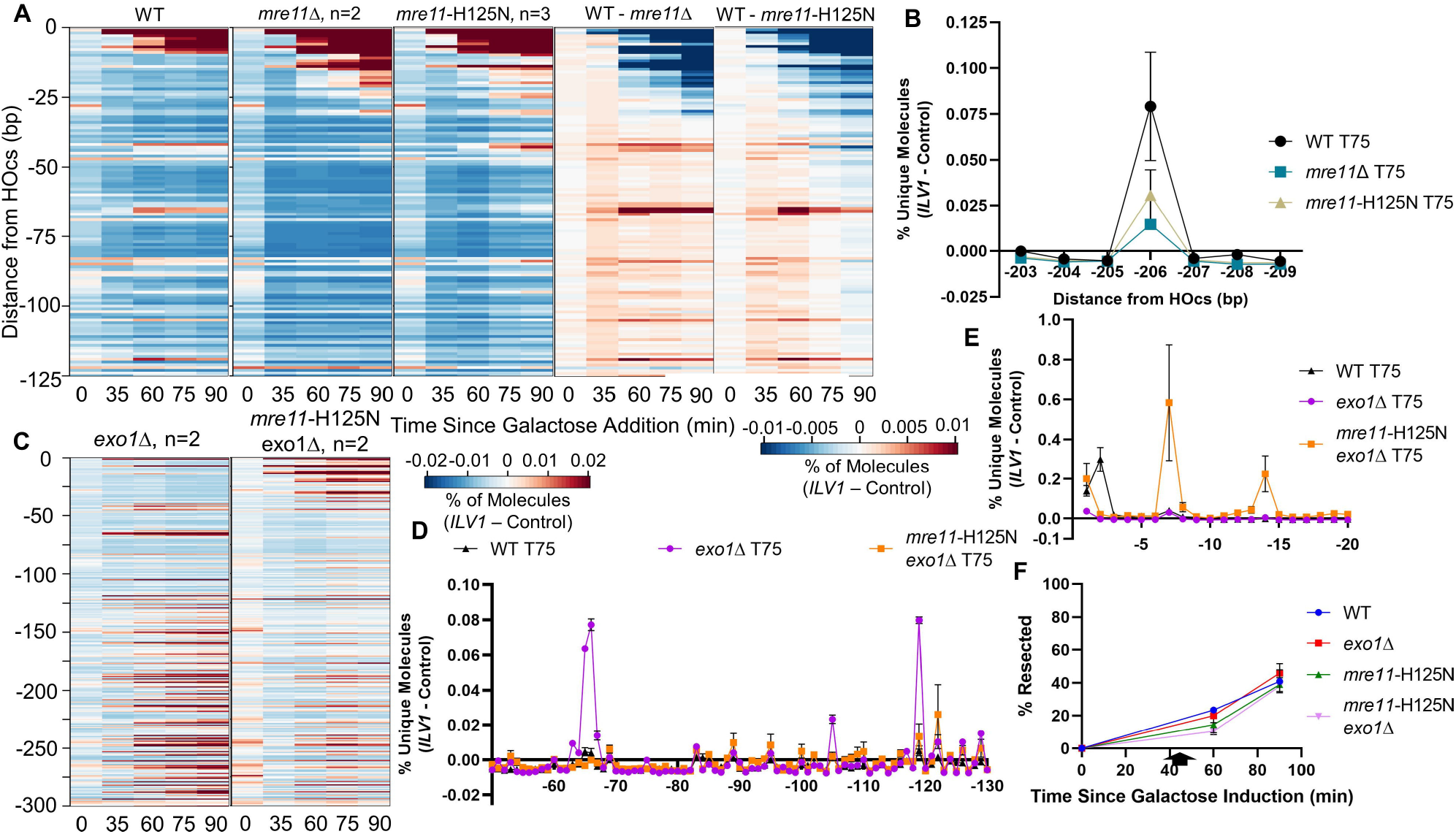
Mre11 nuclease activity is necessary to form resection intermediates at −65/-66 bp. **(A)** The first three panels show heatmaps of resection signal for WT, *mre11*Δ and catalytically defective *mre11*-H125N strains in the first 125 bp from the HOcs. The next two panels plot the difference between the WT and mutant strains to highlight where signal was higher (red) or lower (blue) in WT as compared to mutant. **(B)** Fractional resection signal at positions −203 to −209 bp in WT, *mre11*Δ and *mre11*-H125N strains at the 75 minute time point. **(C)** Heatmaps of resection signal for *exo1*Δ and *mre11*-H125N *exo1*Δ yeast up to 300 bp from the HOcs. **(D&E)** Fractional resection signal 75 minutes after DSB induction in WT, *exo1*Δ and *mre11*-H125N *exo1*Δ yeast at positions **(D)** −50 to −130 bp and **(E)** −1 to −20 bp from the HOcs. **(F)** ddPCR-based resection analysis of WT and mutant strains at a site 355 bp from the HOcs. Points are the average of two biological replicates.

To establish the actions of Mre11 more definitively we added the *mre11*-H125N mutation, which abolishes the Mre11 nuclease activity (Moreau et al., 1999), to our WT and *exo1*Δ strains. The *mre11*-H125N strain by itself behaved similarly to *mre11*Δ implicating the Mre11 nuclease as being required for the increase in distal resection intermediates (Figure 3A). Examination of *mre11*-H125N *exo1*Δ relative to *exo1*Δ revealed a complete loss of signal at −65/-66 and substantially reduced signal at positions −119 and beyond (Figure 3C,D). These results expose an additional stepwise cleavage event mediated by Mre11 at position −119, 53 bp from position −66. Notably, at both the −65/-66 and −119 positions the signal peaked in *EXO1* wild-type strains at 60 minutes and then decreased whereas it remained persistent in *exo1* mutant strains (Supplementary Figure 5), consistent with a kinetic model in which Exo1 acts later at the activated nick created by Mre11. The −119 position appears to be a major transition point as reads are suppressed in the region between −65 and −119 and increased after −119 through the end of our sequencing window in *exo1* mutant yeast.

In contrast, we observed a marked increase in intermediates mapping near the HOcs in the *mre11*-H125N *exo1*Δ double mutant (Figures 3C,E). Resection mediated by Sgs1-Dna2 is the primary pathway in the absence of both Mre11 nuclease activity and Exo1 (Mimitou and Symington, 2008). Therefore, the increase in reads near the DSB end in *mre11-*H125N *exo1*Δ might reflect a shift to resection carried out from the end by Sgs1-Dna2 as opposed to resection beginning from the nick created by Mre11. Intriguingly, analysis of resection kinetics through our sequencing window via the RE-ddPCR assay showed only a small resection defect in *mre11*-H125N *exo1*Δ double mutant yeast compared to *mre11*-H125N and *exo1*Δ single mutants alone (Figure 3F). There is strong evidence that Mre11 has end-chewing activity that aids in Sgs1 loading (Cejka et al., 2010), but the presence of these intermediates in *mre11*-H125N *exo1*Δ indicates that another protein also has such activity. Data obtained from ddPCR resection measurements suggests that Sgs1-Dna2 mediated resection initiation is only slightly less efficient in the absence of Exo1, Mre11 nuclease activity or both.

### Combined loss of Sgs1 and Exo1 reveals the profile of Mre11 activity permissible at a DSB

It is known that Mre11 is responsible for ∼300 bp of resection in *exo1*Δ *sgs1*Δ double mutant yeast defective in long range resection (Gobbini et al., 2020). Indeed, ddPCR quantification of ssDNA produced from resection in *exo1*Δ *sgs1*Δ yeast showed a decrease in resection efficiency in this strain as compared to WT (Figure 4A). Analysis of *exo1*Δ *sgs1*Δ yeast in our sequencing assay revealed an increase in signal at similar positions as observed in the single *exo1*Δ mutant strain (Figures 4B,D, note the scale difference in Figure 4D). The only locations where reads were decreased in *exo1*Δ *sgs1*Δ compared to *exo1*Δ or *sgs1*Δ single mutants were at the positions immediately proximal to the DSB termini, indicating that end chewing events in resection are dependent on the presence of Sgs1 or Exo1.

**Figure 4.**
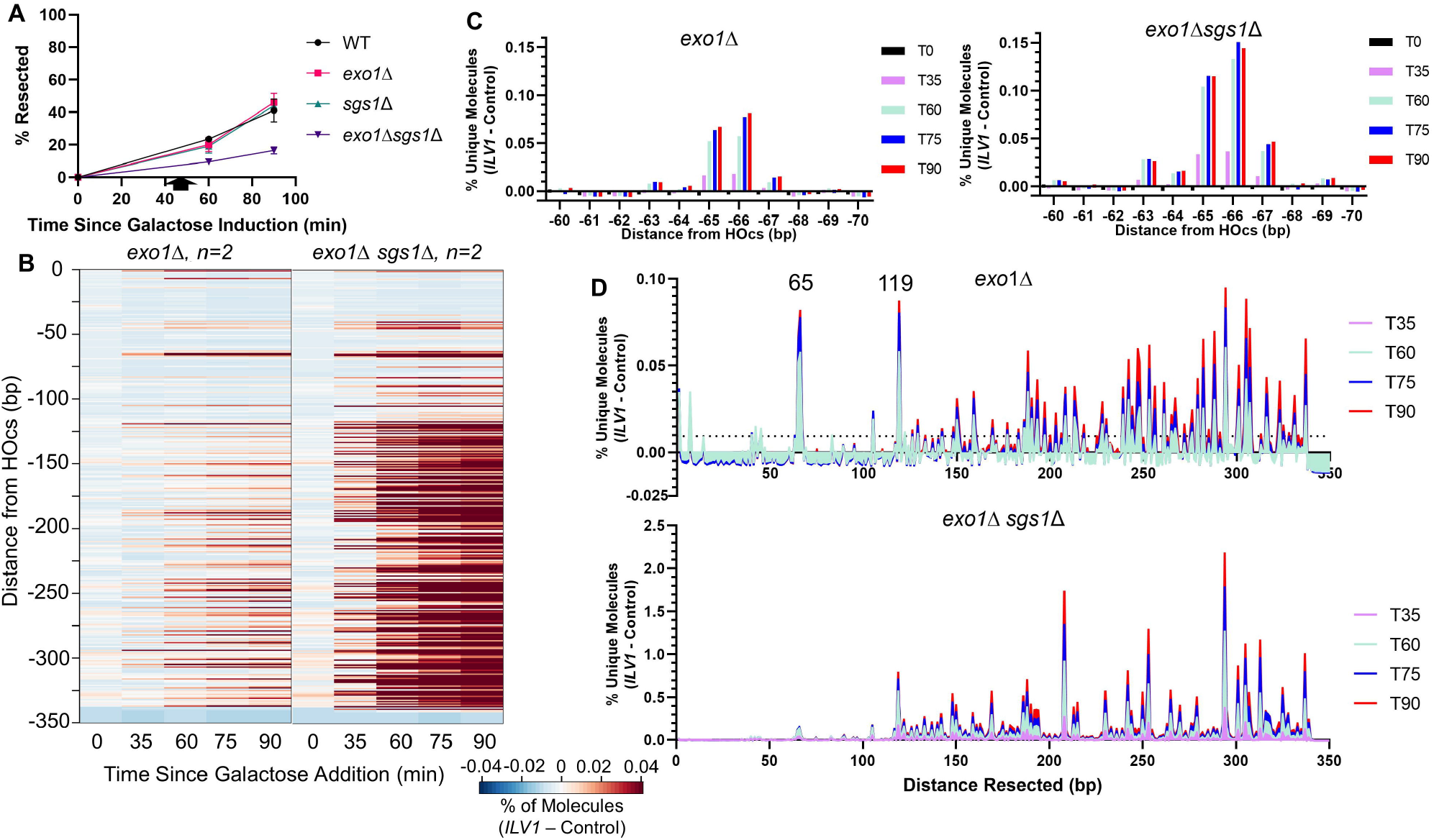
Exo1- and Sgs1-deficient yeast show greatly enhanced resection intermediates beyond position −119. **(A**) ddPCR-based resection analysis of WT and mutant strains at a site 355 bp from the HOcs with combinations of *exo1* and *sgs1* deletions. **(B)** Heatmaps of resection signal for *exo1*Δ and *exo1*Δ *sgs1*Δ yeast. The uniquely mappable *ILV1* sequences end at position −344. **(C)** Fractional resection signal surrounding the −65/-66 bp position for *exo1*Δ and *exo1*Δ *sgs1*Δ yeast. The same y-axis scaling is used on both graphs. **(D)** Fractional resection signal for the first 300 bp away from the DSB for each time point in *exo1*Δ and *exo1*Δ *sgs1*Δ yeast. Note the large difference in y-axis scaling due to the accumulation of many more intermediates in the *exo1*Δ *sgs1*Δ strain.

Interestingly, the percentage of unique reads at the −65/-66 bp position was only modestly elevated in *exo1*Δ *sgs1*Δ as compared to *exo1*Δ (Figure 4C), which contrasts with the large (>10-fold) increase in reads throughout the sequencing window further from the DSB in the double mutant (Figure 4B,D). We observed intermediates mapping to positions throughout our sequencing window without a drop off in signal, which indicates that Mre11 or another protein is resecting past the positions that can be mapped by the assay (344 bp), even when we extended our window to include reads mapped to the common sequence shared with the control allele (Supplementary Figure 6). The massive increase in signal in *exo1*Δ *sgs1*Δ clearly exposed the transition point at position −119 (Figure 4B,D). DSB-proximal to this point, the presumptive actions of Mre11 are restricted to only a few selected positions that have an imperfect periodicity (Supplementary Figure 5). After −119 bp, the resection pattern shifts in both *exo1*Δ *sgs1*Δ and *exo1*Δ to one indicative of a kinetically slower process, since once again cleavage at the more distal positions would prevent detection of simultaneous cleavage at the more proximal positions. It is noteworthy that major peaks occur at positions −208 and −294 in the double mutant, which may represent further stepwise incisions generated by Mre11.

### Mre11 endonucleolytic activity is influenced by DNA sequence preferences

Broadly, three mechanisms might dictate the specific resection profiles seen above: (i) proteins might “measure” cleavage positions relative to the DSB end, (ii) pre-damage factors such as nucleosomes or other bound proteins might constrain repair protein action, or (iii) repair proteins themselves might show a driving sequence dependence. To distinguish these in our system, we made two deletions near the −65/-66 position in WT and *exo1*Δ backgrounds (Figure 5A,C). The deletions each removed three base pairs either 31-33 or 71-73 bp away from the HOcs, i.e. DSB proximal and distal to the −65/-66 position, respectively (Supplementary Figure 1B). Strikingly, the DSB-proximal deletion led to a −3 bp shift in the position of the Mre11-dependent peak in both WT and *exo1*Δ cells, meaning that the same underlying sequence was being cleaved by the Mre11 nuclease even though it was now closer to the DSB end (Figure 5B). Somewhat unexpectedly, the deletion distal to the −65/-66 position changed the pattern of peaks at positions −60 to −66, likely due to their proximity to the deletion (Figure 5D). Both deletions further shifted the resection pattern throughout the distal portions of the sequencing window, including the Exo1-dependent −206 bp position in WT cells and the −119 position in *exo1*Δ (Supplementary Figure 7). Thus, the signal patterns throughout the interrogation window proved to be highly sequence, not distance, dependent.

**Figure 5.**
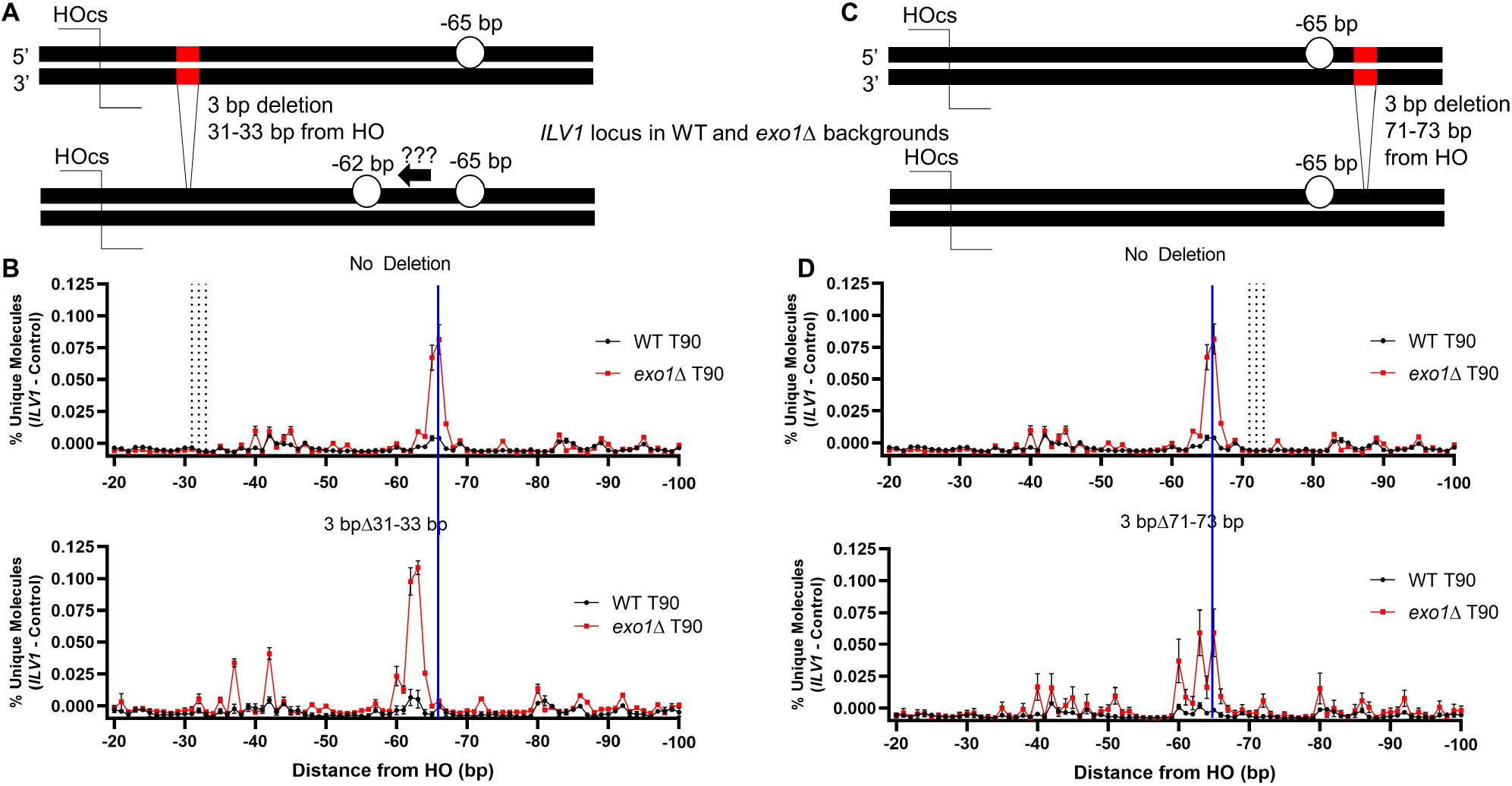
The predominant Mre11-dependent incision shows sequence dependence more than a strict positional dependence. **(A)** Diagram of a 3 bp deletion in *ILV1* located 31-33 bp from the HOcs. **(B)** Fractional resection signal from positions −20 to −100 in WT and *exo1*Δ yeast 90 minutes after DSB induction with and without the 3 bp deletion at positions −31 to −33. Vertical dashed lines in the top plot mark the location of the positions that were deleted in the bottom plot. A vertical blue line indicates the position expected for the peak in the bottom plot if the distance from the DSB end had been maintained after the 3 bp deletion. **(C)** Diagram of a 3 bp deletion in *ILV1* located 71-73 bp from the HOcs. **(D)** Similar to (B) for the 3 bp deletion located at positions −71 to −73.

### Sequencing of 5’ endpoints formed from both ends of the same DSB

To expand our base of observation we next examined the opposite side of the *ILV1* DSB using the same conceptual approach as in Figure 1 (Supplementary Figure 1A). Because this 2^nd^ side of the DSB corresponds to the *ILV1* coding region, we refer to it as *ILV1*-CR and the sequence studied above as *ILV1*-PR, for “promoter” (Figure 6A). We observed a higher background in WT and *exo1*Δ cells for *ILV1*-CR, but a similar overall pattern was observed on either side of the break (Figure 6B). Intriguingly, a number of reads lined up at the *ILV1*-CR −206 bp position in both WT and *exo1*Δ cells. Of primary note was that a peak appeared at −70 bp for *ILV1*-CR and not at −65/-66 bp as for *ILV1*-PR, consistent with the sequence, not distance, dependence noted above. Indeed, analysis of the underlying sequence, oriented similarly with respect to the DSB, revealed a similar palindromic “TCT” motif, with the cytosine being an efficient incision point (Figures 6C,D). Examination of the region around −119 in *ILV1*-CR *exo1* mutants revealed read accumulations at −125 and −153, 55 and 83 bp after the −70 position respectively.

**Figure 6:**
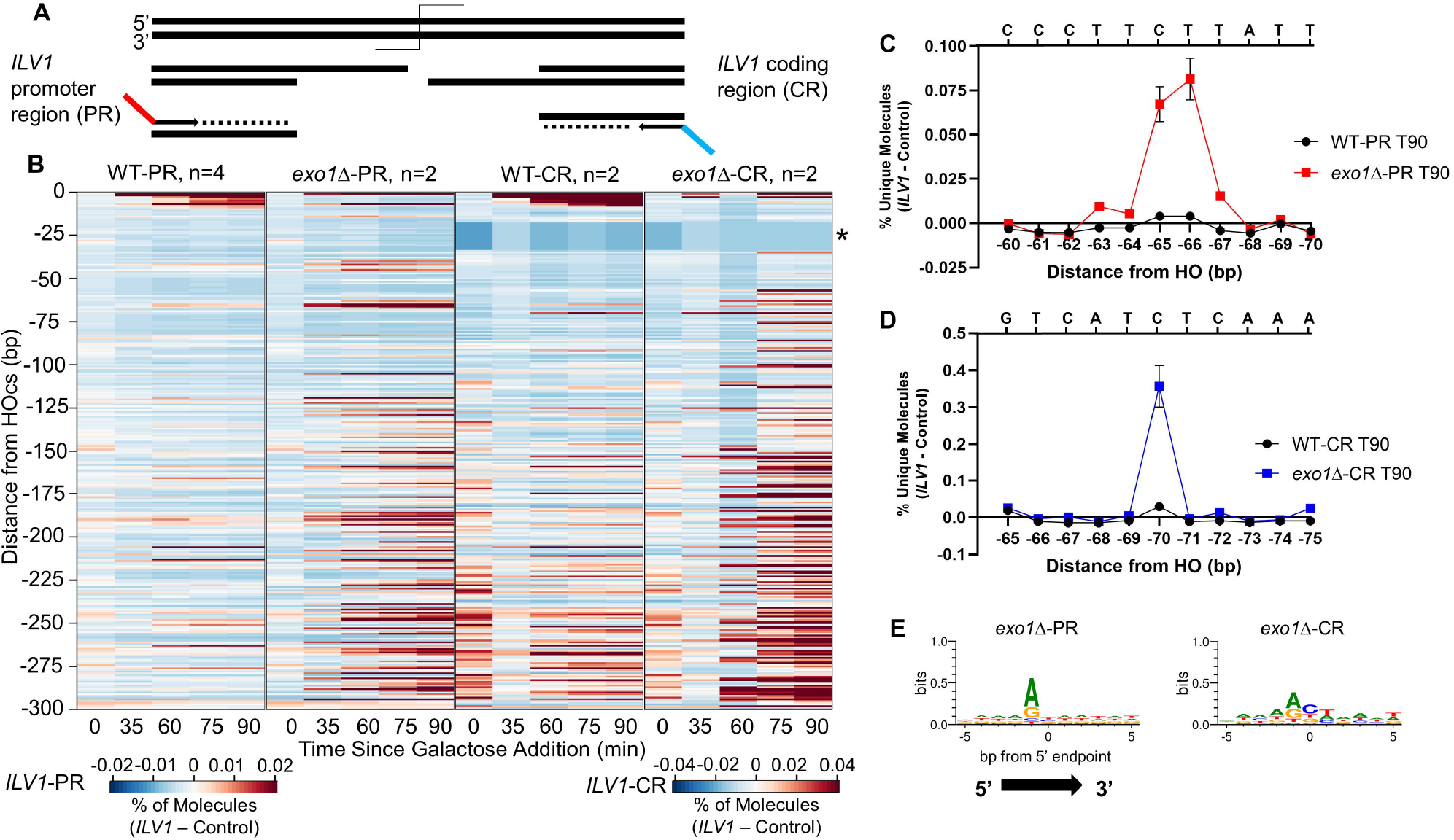
Comparing two sides of the *ILV1* DSB reinforces sequence dependence more than a strict positional dependence. **(A)** Sequencing scheme to capture intermediates resecting into the *ILV1* coding region (*ILV1*-CR; Figures 1-5 examined resection into the *ILV1* promoter region, *ILV1*-PR). See annotated map and sequences in Supplementary Figure 1. **(B)** Heatmaps of resection signal up to 300 bp from the HOcs in WT and *exo1*Δ strains sequenced from either *ILV1*-PR or *ILV1*-CR in separate libraries. The scale for *ILV1*-CR strains is 2-fold less sensitive due to higher background levels. The *ILV1*-CR region marked with an asterisk failed to return any aligned reads for technical reasons due to an adjacent T-rich stretch of bases. **(C)** Fractional resection signal at positions −60 to −70 bp in WT and *exo1*Δ yeast sequenced from *ILV1*-PR. The sequence of the 5’ terminated strand is shown on the upper X-axis. **(D)** Fractional resection signal at positions −65 to −75 in WT and *exo1*Δ yeast sequenced from the *ILV1*-CR, with the sequence as in (C). **(E)** Logo plot analyzing sequences +/-5 bp in either direction from signal peaks in *exo1*Δ yeast sequenced from either *ILV1*-PR or *ILV1*-CR. A threshold of 10% of the max peak height was used to determine the plotted peaks: *ILV1*-PR n=75, *ILV1*-CR n=67. See Supplementary Figure 8 for depiction of these thresholds relative to peak intensities.

To examine the nucleotides contributing to the *exo1*-dependent rise in resection signal in both sequencing windows, we first set a cut-off value of 10% or 20% of the highest number of unique reads counted at one position in each *exo1*Δ mutant. We then generated logo plots of the DNA sequence surrounding these 5’ terminated endpoints (Figure 6E, Supplementary Figure 8). We observed a slight preference for A or G in the position immediately before the 5’-terminated endpoint in Exo1-deficient cells. Even though this sequence-specific signal was weak, it is noteworthy because it is different that the putative motif noted above and thus suggests a non-Mre11-dependent mechanism for many signal peaks. However, a C preference on just the *ILV1*-CR side of the DSB became apparent for the more restricted set of peaks at the 20% threshold (Supplementary Figure 8), which might match the possible Mre11 motif noted above.

### NHEJ and other repair factors influence resection initiation near the DSB terminus

To gain further insight into the obstacles that Mre11 faces near the DSB terminus, we deleted the NHEJ genes *YKU70* and *NEJ1* and analyzed early resection sequencing intermediates. The signal at position −65/-66 persisted in *yku70*Δ, indicating that Ku binding to a DSB end is not uniquely responsible for this Mre11-dependent cleavage position (Figure 7A). However, the more proximal −42 bp peak became more prominent in *yku70*Δ. Moreover, loss of Ku created an interesting change near the DSB wherein WT showed a progressive increase in signal over time whereas *yku70*Δ had a nearly constant signal (Figure 7B), which might suggest an easier movement of resection inward from the DSB end. More drastic was the decrease in reads aligning near the HOcs in *nej1*Δ, which may also signify an increase in the kinetics of resection initiation. Restriction-based ddPCR measurements of end resection kinetics confirmed a higher occurrence of ssDNA at a site 355 bp from the HOcs in these mutants compared to WT (Supplementary Figure 9).

**Figure 7:**
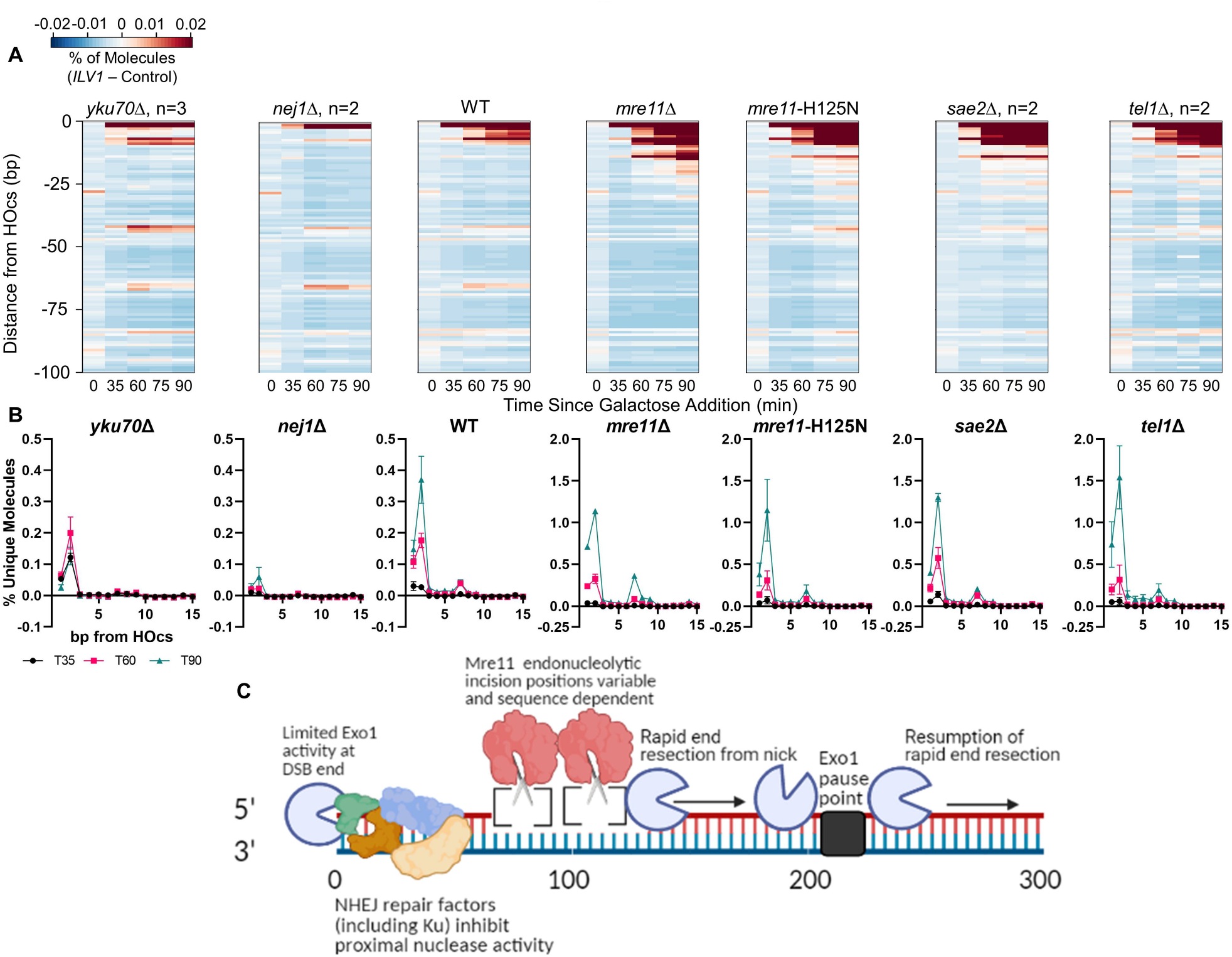
NHEJ factors limit processing events near the DSB end. (**A**) Heatmaps of resection signal within 100 bp of the *ILV1*-PR HOcs in WT and various single deletion mutants. **(B)** Fractional resection signal within the first 15 bp after the *ILV1*-PR HOcs for the same mutants as in (A). **(C)** Model for the initiation of mitotic end resection in yeast that expands on prevailing bi-directional and stepwise models as informed by single base resolution resection monitoring. See text for discussion.

Finally, examination of factors involved in supporting cleavage by Mre11 revealed an increase in reads aligning near the HOcs in *sae2*Δ and *tel1*Δ (note the scale difference between WT and these mutants in Figure 7B) and the same near absence of reads at −65/-66 bp as in *mre11*Δ mutants. These results confirm a slower 5’ degradation process from the DSB end in the absence of Mre11 nuclease activity, specifically at the Exo1-dependent −2 position.

## Discussion

End resection is a fast process occurring at a rate of 4 kb/h (Zhu et al., 2008). However, HR must overcome multiple obstacles close to the DSB that dictate the successful transition to long-range resection (Marini et al., 2019). One such obstacle is repair by NHEJ. We exploited catalytically defective Dnl4 to remove this competing outcome while maintaining normal Ku and DNA ligase IV assembly at DSB ends (Chiruvella et al., 2013). Despite these obstacles, >20% of HO-cut *ILV1* alleles were resected past our ∼350 bp sequencing window within 60 minutes of suffering the DSB. Resection steadily increased to ∼40% of alleles at 90 minutes but was associated with only limited detectable 5’-endpoint patterns in strains that were wild-type for resection-associated genes. Thus, while resection was clearly initiating in the sequenced regions during the experiment, the transition to progressive long-range resection was normally kinetically very fast.

Mutations further revealed important features of the observed patterns, summarized in Figure 7C, where interpretations are guided by the following principles. When a signal peak is observed, we infer that there is a relative delay in processing at that base position. When a strain alteration causes such a signal to increase then that delay has been accentuated, which implicates the altered protein as being responsible for normal processing away from that 5’-terminated intermediate. In contrast, decreased signal in response to a strain alteration suggests that the protein factor normally creates the signal; such an intermediate can sometimes be inferred to be the product of a nuclease’s action. However, a caveat is that primer extension only reveals the most DSB-distal 5’ cleavage point on a DNA strand, so that DSB-proximal increases could result from decreased endonucleolytic activity more distally and vice versa.

Unlike our base *dnl4*-K466A mutation, *yku70* and *nej1* mutations removed the Ku and Nej1 NHEJ factors thought to be inhibitory to resection (Chanut et al., 2016; Sorenson et al., 2017). The greatest effect of these losses was indeed to decrease the signal near the DSB. Thus, Ku and Nej1 might normally impede further processing away from these positions. Loss of Nej1 had the largest impact and reduced signals to near-background levels, consistent with its function in limiting end processing that initiates resection (Mojumdar et al., 2019). An alternative interpretation is that the DSB-proximal signals are not resection intermediates but instead reflect processing associated with futile attempts at NHEJ. These are not exclusive concepts and the fact that signal at the prominent −2 position failed to accumulate in Ku mutant yeast suggests that NHEJ factors might normally, but only temporarily, stabilize DSB-proximal intermediates.

Notably, signal at the −2 position was uniquely dependent on Exo1, which identifies Exo1 as the nuclease that cleaves some, but not all, of the DSB-proximal 5’ intermediates (Gobbini et al., 2018). In contrast to Ku and Nej1, loss of Mre11 or its nuclease activity led to increased signal at the DSB-proximal positions, most profoundly at −2 bp, with a similar result upon loss of the Tel1 and Sae2 proteins that support Mre11 (Cannavo and Cejka, 2014; Cassani et al., 2019; Yu et al., 2019). Here, the best interpretation is that the signal increase is secondary to decreased DSB-distal endonucleolytic cleavage.

Moving inward from the DSB end we encounter the most critical region for resection initiation. Previous *in vitro* analysis of Mre11 activity on Ku-occluded dsDNA substrates revealed incisions at 35-45 and 55-65 bp from the end on 70 and 100 bp substrates, respectively (Reginato et al., 2017; Wang et al., 2017), leading to the prevailing model wherein Mre11 creates a nick at a distance enforced by the Ku protein block (Wang et al., 2018). We observed a pattern of iterative peaks throughout this region consistent with the model and with the inference of a “stepwise” cleavage at somewhat, although not precisely, regularly spaced positions (Cannavo et al., 2019). Although subtle, the pattern of multiple *ILV1*-PR peaks at −42, −65/-66, −84, −105 and −119 bp positions were evident in wild-type yeast and enhanced upon loss of Exo1. That increase was strongly dependent on the Mre11 nuclease as evidenced by absence of −65/-66 and −119 signal in an *exo1 mre11*-H125N double mutant. The relative weaknesses of the −42 signal could be secondary to cleavage at −65/-66, but −84 had substantially less activity, establishing that the different cleavage positions are utilized with different efficiencies.

A central question is what determines the pattern of cleavage at mitotic DSBs *in vivo*. Models cited above imply that Ku and/or Mre11 protein sizes are critical factors (Reginato et al., 2017; Wang et al., 2017; Wang et al., 2018), but the pattern could be enforced by other proteins bound prior to damage such as histones or other proteins (Li et al., 2020; Yu et al., 2018) and DNA repair proteins can have sequence-specific properties (Rahal et al., 2010; Tadi et al., 2016). We addressed these issues using 3 bp deletions in the *ILV1*-PR sequence. Strikingly, the −65/-66 peak moved closer to the DSB with the DSB-proximal deletion and thus tracked the local sequence, not the distance from the DSB end. Further evidence for sequence specificity comes from the identification of a similar incision position at −70 bp on the *ILV1*-CR side of the DSB whose sequence shared 5 out of 7 bases with the *ILV1*-PR −65/-66 peak. Critically, this predominant cleavage position did not move in *yku70* mutant yeast but the −42 position did become the most prominent. Thus, Ku itself is not the primary factor leading to the position of the inferred Mre11 incisions but that it may impede access to the more DSB-proximal positions.

It is equally important to consider where resection intermediates were not observed, which was especially noticeable in *exo1 sgs1*, a strain deficient in long range resection thought to mainly support cleavage by Mre11 (Gobbini et al., 2020). Resection initiation signal with this strain increased dramatically, although the signal at −65/-66 was only modestly increased relative to *exo1*. Indeed, most positions up to −119 bp had limited read counts despite the poor ability to transition to progressive resection. We infer that the first ∼120 bp of a DSB are a unique zone where a specific protein “footprint” protects the DNA from promiscuous activity and where Mre11-dependent cleavage is preferred. The source of that protection is not entirely clear, but it is not Ku, Sgs1, or Exo1 based on our data. It might be Mre11-Rad50 itself, although this is difficult to prove with resection data alone.

The genesis of signal after −119 bp is intriguing. In yeast with Exo1 and Sgs1 it is dependent on the Mre11 nuclease. This is especially true for the strong and specific −206 position, which we believe results from exonucleolytic extension away from the more proximal incisions. Consistently, the −206 signal was abrogated in *exo1* yeast suggesting it as a pause point during extension by Exo1. We do not know the nature of the obstacle that causes Exo1 to pause but one candidate is the 9-1-1 complex based on recent observations (Gobbini et al., 2020). Whatever is responsible is again sequence specific as the location of the peak shifted with the DSB-proximal 3 bp deletions. The signal accumulation at nearly every position past −119 bp in *exo1 sgs1* yeast is uncertain; it might be a result of more distal, kinetically slower and pathologic cleavage by Mre11 but we were unable to make an *exo1 sgs1 mre11*-H125N triple mutant to test this idea (Mimitou and Symington, 2008).

A caveat is the possibility for artifacts arising from the use of the HO endonuclease. HO can remain bound to its cleavage product (Jin et al., 1997) so we cannot rule out that HO itself is partially responsible for the observed cleavage patterns, although all DSB repair actions occur at HO DSBs and we make our measurements long after repair proteins have engaged the DSB. We also considered whether HO might cleave off-target to create some measured endpoints, especially when the DSB was resistant to resection (Nickoloff et al., 1990). Importantly, we never saw anything above a low baseline cleavage at the control allele. Also, many cleavage intermediates could be abrogated by repair mutations identifying them as specific. However, one peak at –294 in *exo1 sgs1* yeast had possible sequence similarity to the HOcs and could represent “skipping” of HO to this alternative position (Supplementary Figure 10).

Our results align with studies that used sequencing approaches to study meiotic end resection. Notably, we observed increased intermediates throughout our sequencing window in *exo1* cells that mirrors the effect reported by the Keeney group (Mimitou and Keeney, 2018; Mimitou et al., 2017). Our data also support the observations that Sae2 and Tel1 promote efficient resection and a decrease in unique reads mapping near the DSB termini (Mimitou and Keeney, 2018; Mimitou et al., 2017; Paiano et al., 2020). Keeney described evidence of Exo1-dependent pausing at heterochromatin but it is uncertain how closely that phenomenon corresponds to our precise Exo1 pause point (Mimitou et al., 2017).

In summary, results here provide a uniquely high-resolution picture of mitotic DSB resection initiation. They support published models in which Ku and Nej1 inhibit exonucleolytic resection from the DSB end in a manner that is overcome by Mre11 incision with a peak activity ∼60 to 70 bp from the DSB end but with evidence for multiple incision points that might occur in a stepwise fashion. Subsequent nick extension by predominantly Exo1 is rapid and processive but also subject to pausing (Figure 7C). However, the cleavage pattern is complex, substantially sequence dependent, and influenced by an inferred protective protein complex that is not strongly dependent on Ku. Critical challenges are to extend these findings across many DSBs and to correlate DNA-based results with equally high-resolution protein binding studies *in vivo* to understand the nature of the early mitotic repair protein complex.

## Materials and Methods

### Yeast strains, growth media and DSB allele

Haploid yeast were grown at 30 C in either rich medium containing 1% yeast extract, 2% peptone and 40 μg/ml adenine (YPA) with either 2% glucose, 2% galactose, or 3% glycerol as a carbon source. Yeast strains were isogenic derivatives of BY4741 (Brachmann et al., 1998). The base strain for most strains used in this study was YW3104, itself a derivative of YW1405 that has been previously described (Wu et al., 2008). All additional gene disruptions and modified alleles were made using a PCR-mediated technique or a *URA3* pop-in/pop-out method and confirmed by PCR and sequencing (Palmbos et al., 2005). A table of all yeast strains is provided as Supplementary Table 1.

Relative to YW1405, YW3104 bears a previously described point mutation, *dnl4*-K466A, that abrogates activity of the yeast NHEJ ligase (Chiruvella et al., 2013). A further change at the *ILV1* locus was made to introduce the NdeI site and other sequences used in most experiments; its exact content was related to a prior design concept that is not discussed here. Specifically, we inserted a bacterial *ori* sequence followed downstream by the *HIS3-MX6* selection marker at a genomic location 802 bp upstream of the *ILV1* Hocs (Supplementary Figure 1). The fourth change in all strains that were subjected to sequencing-based resection monitoring was the addition of a control allele, which was always the last construction step to yield paired strains with and without the control allele. For this purpose, we identified an appropriate native RE site in the genome (e.g. NdeI) and inserted a construct containing homology to *ILV1* 350 bp upstream of the RE cleavage site with *URA3* as a selection marker. This construct lacked the HOcs but was homologous to the portion of *ILV1* that is needed for primer extension and library amplification (Supplementary Figure 1).

### Oligonucleotide primers and probes

Oligonucleotide primers and probes were from IDT. A complete list of all sequences is provided in Supplementary Table 2. All primers bore 5’ hydroxyls. Primer extension primers were made with hand mixed UMIs at an equal percentage of A,C,G,T and a series of 6 different known and fixed time point barcodes. They were designed to anneal to the RE site DBS-distal to the HOcs such that primer extension reaction would copy both the 5’ primer tail (from the genomic DNA 3’ end) and the genomic DNA (from the primer 3’ end). Replicate time courses from the same strain always used different primer extension primers to ensure that the time point barcodes did not create an unanticipated bias.

### DSB induction paradigm

Strains containing the *ILV1* HOcs with and without the control allele were grown separately overnight at 30 C to saturation in YPA-dextrose media then inoculated to a low OD_600_ in YPA-glycerol for a consecutive night. When the OD_600_ of the YPA-glycerol cultures was in the range of 0.3 to 0.6, the strain with the control allele was spiked into the other YPA-glycerol culture at a ratio of 1:20 as determined by OD_600_. The T0 time point was taken by rapidly mixing 45 mL of this YPA-glycerol culture into 5 mL 0.5 M EDTA to quickly quench cellular nuclease activities. Galactose was then added to the YPA-glycerol culture at 2% final concentration to induce HO expression and DSB formation. Another time point was taken 35 minutes after galactose addition in the same fashion as T0. At 45 minutes after galactose addition the YPA-glycerol+galactose culture was pelleted and the cells resuspended in an equal volume of YPA-dextrose followed by continued shaking at 30 C. Further time points were taken as above at 60, 75 and 90 minutes after galactose addition (15, 30 and 45 minutes after the switch to dextrose, respectively). Time course materials used in ddPCR-based resection monitoring were generated by a similar protocol but without the control allele spike-in and only 1.5 mL of cellular extract was taken from a smaller total growing culture volume without EDTA addition.

### Genomic DNA preparation and quality validation

High molecular weight gDNA was obtained using the Thermo Scientific Pierce Yeast DNA Extraction Kit. Yeast cells were thus lysed without glass beads or damaging chemical treatment. Genomic DNA was subjected to 0.8% agarose gel electrophoresis to confirm its high molecular weight and purity (Supplementary Figure 3). Importantly, DNA is less prone to random fragmentation near the end of a DNA fragment, which was true for all stretches of DNA we studied except for the ILV-CR side of the DSB at time 0 before DSB formation. Those molecules placed the interrogation region in the middle of a large RE fragment, which resulted in notably higher background as compared to the control allele at T0 (Figure 6A). As expected, this phenomenon resolved by T35 because the HO DSB now placed the interrogation region at the end of the source DNA molecule.

### Resection monitoring by RE-ddPCR

To measure resection flux through the interrogation window of our sequencing method, we used a ddPCR method based on resistance of ssDNA to RE cleavage. That method is described in detail in Lomonaco et al. (Lomonaco et al., 2021). Briefly, HinfI was used to digest 300 ng gDNA from the smaller 1.5 mL cultures. HinfI cleavage was confirmed by gel electrophoresis. ddPCR measurements using probes targeting a HinfI site 355 bp from the HOcs and an *ACT1* control locus revealed when the HinfI site became resistant to cleavage due to resection.

### Resection monitoring by high resolution primer extension sequencing

#### Quantification of DNA amount and the exact control allele ratio

gDNA was quantified and HOcs curves were established using a dual fluorophore ddPCR assay with primers flanking the HOcs (FAM) and within an undamaged reference gene, *ACT1* (VIC). The fraction of uncut alleles was directly measured as the fraction of molecules giving signal with the HOcs relative to the *ACT1* probes. The true control allele spike-in percentage was further established in the sequencing assay using DNA extracted from the T0 sample, i.e. prior to DSB formation. That assay used paired probes homologous to the control allele (only present in the spiked-in cells) and the HOcs-containing *ILV1* locus (present in all cells regardless of whether they had the control allele, i.e. all cells in the culture made the same DSB). Importantly, all further time points were taken from the exact same culture so the value from the T0 specimen applied to the later time points also.

#### Restriction enzyme digestion, primer extension and intramolecular ligation

Twenty micrograms of gDNA were digested by 40 units of NdeI (NEB) in Cutsmart Buffer at 37 C for one hour. The tubes were then incubated at 65 C for 20 minutes to deactivate the NdeI. The digested DNA was purified using AMPure XP Beads and eluted using water in a volume of 60 µL. The primer extension reaction used 1 unit of Phusion DNA polymerase in the provided buffer and a custom primer containing a fixed 6 nt time point barcode sequence, 12 nt hand mixed UMIs and several single fixed nucleotides used to locate the UMIs (see below). Primer extension and tagging of molecules was done separately in each time point with different fixed 6 bp barcode sequences. The uncycled extension reaction was: 98 degrees for 3 minutes, 55 degrees for 3 minutes, 72 degrees for 10 minutes. Twenty µL of the eluate following restriction enzyme digestion and purification was added to three separate PCR tubes for each timepoint during this step, with a final volume of 50 µL in every reaction. The extended molecules from each time point were purified using AMPure XP and eluted using water in a final volume of 40 µL. The entire eluate from each time point was diluted into separate overnight intramolecular ligation reactions at room temperature using T4 DNA Ligase (NEB) at a final concentration of 6,400 units in a 320 µL reaction volume in the provided buffer. The ligase was then inactivated at 65 C for 10 minutes.

#### Amplification and sequencing of custom library

Once UMIs had been covalently ligated to gDNA 5’ endpoints and the ligase inactivated, all time point samples from a given culture’s time course were pooled together and purified into 200 µL water using AMPure XP beads. Therefore, any bias due to PCR amplification and sequencing was expected to apply equally to all time points. Specifically, ligation junctions were amplified with primers flanking the HOcs that were tailed with sequences homologous to Illumina P5 (DSB side) and P7 (upstream NdeI side) adapters used in high-throughput sequencing. This reaction used KAPA HiFi HotStart Polymerase with the following conditions: 95 C for 3 minutes, 17 cycles of 98 C for 20 seconds, 60 C for 15 seconds, 72 C for 30 seconds, and finally 72 C for 1 minute. Twenty µL of the ligated and purified eluate was added to 10 separate PCR tubes, each a final volume of 50 µL, for every pooled strain time course during this amplification. Products were purified on AMPure XP beads and eluted in 125 µL then a custom Illumina amplicon library was made through PCR using IDT Illumina-compatible i5 and i7 indexed adapters (IDT10_UDI_Adapter pairs #97 to #106), which identified the time course, i.e. source sample culture. The second adapter PCR also used KAPA HiFi with the following conditions: 95 C for 3 minutes and 8 cycles of 95 C for 30 seconds, 55 C for 30 seconds, 72 C for 30 seconds. Fifty nanograms of the purified previous amplification product was put into this PCR with a final volume of 50 µL for each library. Ten individual custom amplicon libraries, each representing one complete time course series, were submitted to the University of Michigan Advanced Genomics Core where they were balanced, pooled and sequenced on 2.5% of a shared Illumina Novaseq lane in the 2 x 150 format.

### Bioinformatics analysis

#### Code and file availability

All code and support files required to execute data analysis are available from GitHub at https://github.com/wilsonte-umich/resection-seq. The pipeline was executed using the q pipeline manager available at https://github.com/wilsonte-umich/q-pipeline-manager. Briefly, a q data file establishes values for sample-and server-specific variables, which then invokes repository file ‘resection-seq/process_sample.q’ via command ‘q submit <data-file>.q’. Each DSB end to be profiled required an indexed FASTA file with the sequence of the reference and control alleles, plus Perl and R scripts defining their structures. Files for all DSBs used in this study are provided in repository directory ‘resection-seq/Bazzano_2021/references’. Each sample to be analyzed required a row entry in a metadata file that described the experiments and time points; a complete file for all samples in this study is provided as ‘resection-seq/Bazzano_2021/ResectionMasterStrainTable.txt’.

#### Read validation and alignment

Data from the sequencing core consisted of paired end reads that had been de-multiplexed per sample using the Illumina indices added in the last PCR step but that still had a mixture of all time points per sample. Read 2 (from the P7 primer) was only needed for sample demultiplexing and was not used further. Each read 1 from the P5 primer was analyzed to determine if it matched, in read order, (i) an appropriately long leader sequence corresponding to the DSB-distal end of the resected fragment, (ii) a UMI of the appropriate length, (iii) a single fixed A base, (iv) a known 6 bp time point barcode, (v) two fixed TA bases, and finally (vi) sequence corresponding to the DSB-proximal or control-proximal DNA that had been ligated to the UMI during circularization. Reads that failed to match this pattern were discarded.

For matching reads, 36 bp of the proximal sequence was extracted and aligned to the DSB and control reference sequences using Bowtie (Langmead et al., 2009) with options ‘-v 3 -k 1 -m 1 –best’, such that only unique alignments with up to three mismatches were accepted (some reads were expected to map twice to the common sequence shared between the DSB and control alleles). Productive alignments were sorted and grouped by all unique combinations of time point, UMI sequence, and alignment position, which acted as the definitive molecule identifier (a UMI could be used by more than one time point because those libraries were prepared separately and by more than one mapped position because different positions were distinctive based on the strict alignment criteria). Metrics consistently confirmed high data quality, including that (i) there were <2% unaligned reads, (ii) reads only aligned to the predicted strand of the reference sequences, and (iii) >95% of reads aligned with no mismatches.

#### Purging redundant UMIs

We next accounted for PCR or sequencing errors introduced into UMIs that would lead to over-counting of the number of truly unique source molecules. We used custom code to implement the directional model of Smith et al. (Smith et al., 2017), which considers a UMI network to have originated from the UMI node with the highest counts, with UMIs added to an existing network if they have at most 1 mismatch from a UMI already in the network. Script ‘collapse/collapse_UMIs.R’ implements an efficient algorithm for solving such networks that matched our needs, given that we expected very large numbers of unique molecules mapping to a few specific alignment positions. The algorithm exploits 2-bit encoding of all possible UMI sequences and pre-calculated XOR values for all UMI pairs with a Hamming distance of 1.

Our 12 bp UMIs did not always have sufficient diversity to avoid UMI collisions for the HO or RE-cut ends of the DSB allele; for this reason and for computational speed we disregarded DSB allele positions that had more than 500K mapped molecules, which always included the DSB end itself. In contrast, all positions were always subjected to UMI analysis for the control allele, which had been doped into the yeast pool at a nominal fraction of 5% so that UMI collisions were expected to be minimal. The result was a reduction in molecule counts per unique combination of time point, UMI sequence, and alignment position to reflect only true inferred source molecules.

#### Discarding failed libraries

To identify poorly performant libraries we wrote repository script ‘‘resection-seq/parse_q_report.pl’ to read the output of ‘q report -j all <data.file>.q’ and write a table with summary statistics of all samples. The output is provided for all samples as repository file ‘resection-seq/Bazzano_2021/ResectionOutputSummary.xlsx’. It allowed us to establish criteria that a time point library needed to have at least 3.5M cell equivalents of input DNA, as determined by the corrected control molecule count, and a background breakage frequency less than 0.03% in the control allele. Libraries failing these criteria were written to repository file ‘resection-seq/Bazzano_2021/BadLibraryTable.txt’. Because time point libraries were initially handled separately, a sample could have many successful time points even if one failed.

#### Assembly, normalization and visualization

For each sample we cross-tabulated read counts such that mapped positions were in rows and time points in columns and wrote two R Shiny web tools to analyze these tables, provided in repository directory ‘resection-seq/_server’; ‘visualize’ creates scatterplots for a series of samples as a function of allele position whereas ‘heatmap’ creates plots of position by time point with values expressed by color intensity.

For each tool, the first normalization step was to sum the count of all unique control allele molecules, whether intact or fragmented, to establish the total number of control alleles (and thus haploid cell equivalents) present in the sequenced DNA. We next calculated the inferred number of DSB alleles present in the sequenced DNA by correcting that control allele count by the fraction of control allele-bearing cells in the sequenced mixture, determined by ddPCR from the time 0 specimen as described above. The fraction of all input alleles that mapped to each possible DSB or control allele position was calculated by dividing the molecule count at that position by the actual or inferred allele counts established above. Although degradation of the DSB allele could lead to lower total recovered counts at later time points, the calculated fractions remained accurate because they were normalized to the control allele, which was unaffected by DSB processing.

The signal for the control allele was mostly at the NdeI restriction site (Supplementary Figure 4) but the signal at other control positions provided a means of assessing the extent and nature of random fragmentation that occurred in each DNA prep. This pattern included evidence that smaller PCR fragments were sometimes more highly represented in final libraries, so we characterized the background fragmentation by fitting a linear regression to the non-NdeI positions of the control allele. The fraction predicted by that regression was then subtracted from the fraction values at all DSB positions to establish our final normalized values for DSB allele positions, referred to as “signal above control” or “DSB – control”. Examination of samples at time 0 consistently validated that the control allele regression line was an excellent model for background fragmentation of the DSB allele (Supplementary Figure 4).

Finally, normalized values were averaged for each time point over all samples of the same culture type (e.g. WT). Heatmaps applied a linear color transformation to the averaged DSB signal above control; the scale of all plots is provided in the figures. The value can be negative due to random fluctuations but is never expected to have large negative values.

#### Logo plots

Logo plots were generated using http://weblogo.threeplusone.com/. *ILV1*-PR and CR % GC content (36 and 38%, respectively), were taken into account when generating these plots.

## Supporting information

Supplementary File 1 - library statistics

## Data availability

FASTQ files with sequencing reads are available for all samples from the NIH Sequence Read Archive via accession PRJNA703820 (temporary Submission ID: SUB9120849).

## Acknowledgements

This work was supported by NIH grant R01 GM120767 to T.E.W and a supplement of the same number awarded to S.L. The authors thank Olivia Koues and the staff of the Michigan Advanced Genomics Core for advice and expert handling of high throughput sequencing services. We also thank Christian Rizza for his work assembling the RE-ddPCR-based resection plots and former lab members Dongliang Wu, James Daley and others for input and discussions over the course of developing this project.

