## Supplementary Figures and Tables for "Mapping yeast mitotic 5’ resection at base resolution reveals the sequence and positional dependence of nucleases *in vivo*"

**A**

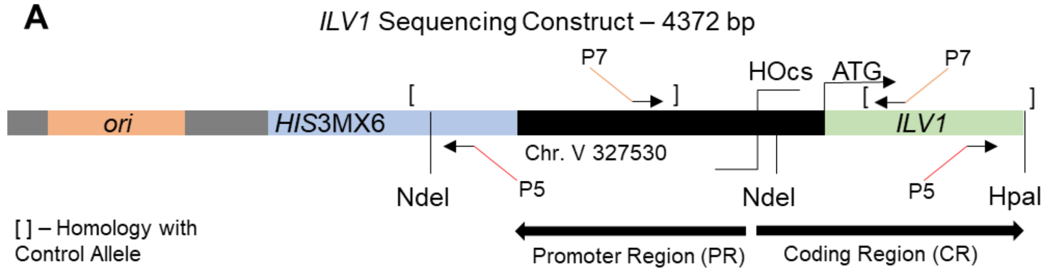


**B** *ILV1*-PR Ligation Junction

```
 P7 -------------------------->
1    AACCTTCCAC ATATCGTCCA ATTCTGCAGC CCACATCTTT TTCCACCACG   50
          End of Homology with Control Allele
51   ATACGGGAAA CAGAATGGGT CCTTGGATTC TCGCTAAACA GTCTCTCTAT 100
101  TCCCCTGTTC AAAACCCTCA AGATATTTGT TTCCCGCAAC AGCTGCAATT 150
151  GCAATTGATC AATCCTATGC GAAAATGCCG AGTTTATGTT ATTCAAGACG 200
                                -206
201  CATTTTAAAA AATTCACTAG CGGCTCCTTG AAATTCATTA TGTCTGATGA 250
251  ATATGAAAAC CTTTTCCTGA CTACCAAGAC TCTTTAACTC TTCTCTCTTT 300
301  ATTGCATATT ATCTCTGCTA TTTTGTGACG TTCAATTTTA ATTGACGCGA 350
            -71    -66,-65                            -31
351  AAAAGAAAAA ATAAGAAGGG CAAAAAGAAA AAGCGCAGCG GGTAGCAAAT 400
                HO Binding       Cut 0
401  TTGGAATCGC TTTTAGTTTC AGCTTTCCGC AACAGTATAA TTTTATAAAC 450
451  CCTGATTTGG AATCGCATAA AAAGAAAAAA AAAATATCAA AGAAAAAGAG 500
                NdeI,
             Uncut 0              Extension Primer
501  TCATCTCAAA CATATAXXXX XXTNNNNNNN NNNNNTGATC CAAAAAGCCA 550
551  ACACCTGTGT GAATATCGAT GACCTGTGAC TGAGTAGCTT GGGAA       595
                <-------------------------------------- P5
```

**C** *ILV1*-PR Control Allele Ligation Junction

```
 P7 -------------------------->
1    AACCTTCCAC ATATCGTCCA ATTCTGCAGC CCACATCTTT TTCCACCACG   50
                End of Homology with ILV1-PR
51   ATACGGGAAA CAGAATGGGT CCTTGGATTC TCGCTAAGTT GTAATTTTGG 100
101  CTAAACCCAG TTCAACATCA CCACAAAAGA AGGTTTCCTT TATGCCCTTA 150
151  CCCTTTCTGC CTTCTTATTC TAGAATTTGT TTTGATGCTG TATTTGTTTG 200
201  GTCCTTGTTC GTCATCGAAA AACCGGCCCA AAAACGGCTA ATAAAAACGT 250
251  GAACCTGTTT TAAGTCTTTT GCTATAAAAG ATATTTATAC TCTAAAAAAA 300
301  GGGAAACTTT TGCAGCAAAA CAAATTTTAT GCATTAGTTC TCTGTTCTTA 350
351  ACTTGGTCAG ACAAAATTCG TCCGTTTGTA GTTGTCTGTA CAGGAATTCA 400
                             NdeI,0      Extension Primer
401  TCGAGCGATA TTCTATCCTG AAATACATAT AXXXXXXTNN NNNNNNNNNN 450
451  TGATCCAAAA AGCCAACACC TGTGTGAATA TCGATGACCT GTGACTGAGT 500
                                  <--------------------------
501  AGCTTGGGAA                                             510
     ---------- P5
```

**D** *ILV1*-CR Ligation Junction

```
 P7 ------------------------->
1   ACACCTTGAG AGATTGGAGA TTCATTAATA ACATCGTATA CAGAGGACCT  50
                   End of Homology with Control Allele
51  TAAAACTAAA CGGACGTAAT CAGGGGTGTT ATCAGTTTGC AGCTCATCCA 100
101 ATTTCAATTC AGAGTGTAGT TTTATCAAGG AAGGTGACAG GTGTTGTCTG 150
151 TGCAAATGAG CCTTTAGTCT CAAAAGGTTC AATCCAGACA CTTTGGACTG 200
                                              -206
201 TTTACCTTGC CGAACAACCG TACATAATGG TTGCTTTAGT AGAGTAGCTG 250
     <-
251 ACATCACTAA CTTTGTGTAA TTGACTTAGT TTAAATGTGG CTTGTAGCAC 300
301 TCACACAAGA AAGAACTCAA AGCAGCAACA ACAAAAGTTT TCAAAGCTGA 350
                                 -70
351 TAATGAAGTA TCTGCAGACA TATGTTTGAG ATGACTCTTT TTCTTTGATA 400
                                    HO Binding      Cut 0
401 TTTTTTTTTT CTTTTTATGC GATTCCAAAT CAGGGTTTAT AAAATTATAC 450
                 Extension Primer
451 TAXXXXXXTN NNNNNNNNNA ACAAGAACAA CTTCATCAAC CACTTGTTGG 500
501 GCGACTCTAA ATGTTTCTTC ACCAATCATA CGCACAGACG TACCAT      546
                                 <---------------------- P5
```

**E** *ILV1*-CR Control Allele Ligation Junction

```
 P7 ------------------------->
1   ACACCTTGAG AGATTGGAGA TTCATTAATA ACATCGTATA CAGAGGACCT  50
                              End of Homology with ILV1-CR
51  TAAAACTAAA CGGACGTAAT CAGGGGTGTT ATCAGTTTGC AGCTCATCCA 100
101 TGTAATAGAT CTAATGAATC CATTTGTTAG TTAATAGTTT AAATGTTTTT 150
151 ATCGGAAGAG GTTTTGTCAT CACATCAGCA ATGTTCTTAT TGGTCTCGAT 200
201 GTAGTATACG TATAAATTAT TACCTGATAC TTCATCTCTA AGTCTCATTG 250
251 CCTTCGTGCC AAAAAATCTG TTTCTAAATT TCTCTTCATT TGTAGACTTA 300
301 ATTATACTGA TCGTTGATCT ACTATCAGTA AGTAAGCCTT TAATAATTGG 350
351 TTTCTTGTTA AGTTCTTGCA CAAGGTGACT GAGGTTATTC AATAGCGGTA 400
                                                    HpaI,0
401 TAGCTTCACT GACTGCGTGT ATTTCTGCTT CTGTAGTTGA AGTGCATGTT 450
                Extension Primer
451 TAXXXXXXTN NNNNNNNNNA ACAAGAACAA CTTCATCAAC CACTTGTTGG 500
501 GCGACTCTAA ATGTTTCTTC ACCAATCATA CGCACAGACG TACCAT      546
                                 <---------------------- P5
```

**Supplementary Figure 1: Map and sequences of the *ILV1* DSB and control alleles.**
**(A)** Map of the *ILV1*-PR and CR target regions. Resection from the HOcs into *ILV1*-PR extended into a non-native sequence inserted 802 bp away at Chr. V position 327530. Restriction enzymes used for *in vitro* isolation of the *ILV1* fragments are shown; note that *ILV1*-CR lacks a HpaI RE site distal to the HOcs and therefore had increased random fragmentation at T0 (see Methods). Brackets denote the homology to the control allele, with 1084 bp of common sequence in *ILV1*-PR and 804 bp in *ILV1*-CR. Primers within the control allele homology region had Illumina P5 and P7 complementary adapter tails and were used to amplify the ligation junction. **(B)** Annotated sequence of ligation junctions at the NdeI site in *ILV1*-PR libraries. Arrows indicate the tailed amplification primers. The homology region shared with the control allele is underlined. The HO recognition sequence is labeled with “Cut 0” referring to the mapped 5’ endpoint position after the DSB was formed. Adjacent are the 4 bp 3’ overhangs left by HO

cleavage (boldface). Distal to this position is the NdeI site (double underline) where reads aligned when the HOcs was intact (Uncut 0). Adjacent (bold underline) is the sequence of the extension primer ligated to the NdeI site. The fixed first, second and seventh bases were used for characterization of sequencing reads. 'X' denotes variable timepoint barcodes whereas 'N' denotes UMI bases. This extension primer sequence moved leftward to a new ligation point upon HO cutting or subsequent resection. Read accumulations discussed in the text and the introduced 3 bp deletions are highlighted. **(C)** The *ILV1*-PR control allele ligation junction, similar to (B). **(D)** The *ILV1*-CR ligation junction. The *ILV1* start codon is highlighted in green. **(E)** The *ILV1*-CR control allele ligation junction.

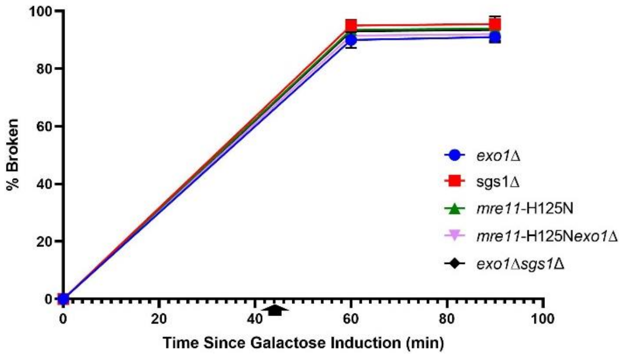


**Supplementary Figure 2: DSB generation is efficient in mutant backgrounds.**
*ILV1* DSB formation efficiency as determined by ddPCR similar to Figure 1A. Like WT, all strains showed >90% DSB formation at the critical time points for resection monitoring.

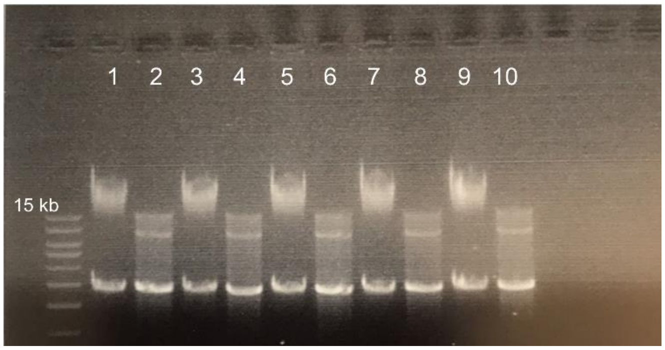


**Supplementary Figure 3: High-molecular weight gDNA used for resection sequencing.**
Genomic DNA extracted in a WT time series was run on a 0.8% agarose gel to determine its molecular weight and purity. The top ladder band is a 15 kb. Odd numbered lanes: ~1 µg of genomic DNA from T0, T35, T60, T75, and T90. Even numbered lanes: the same DNAs digested with NdeI. Note the expected yeast RNA super killer virus band at ~5kb.

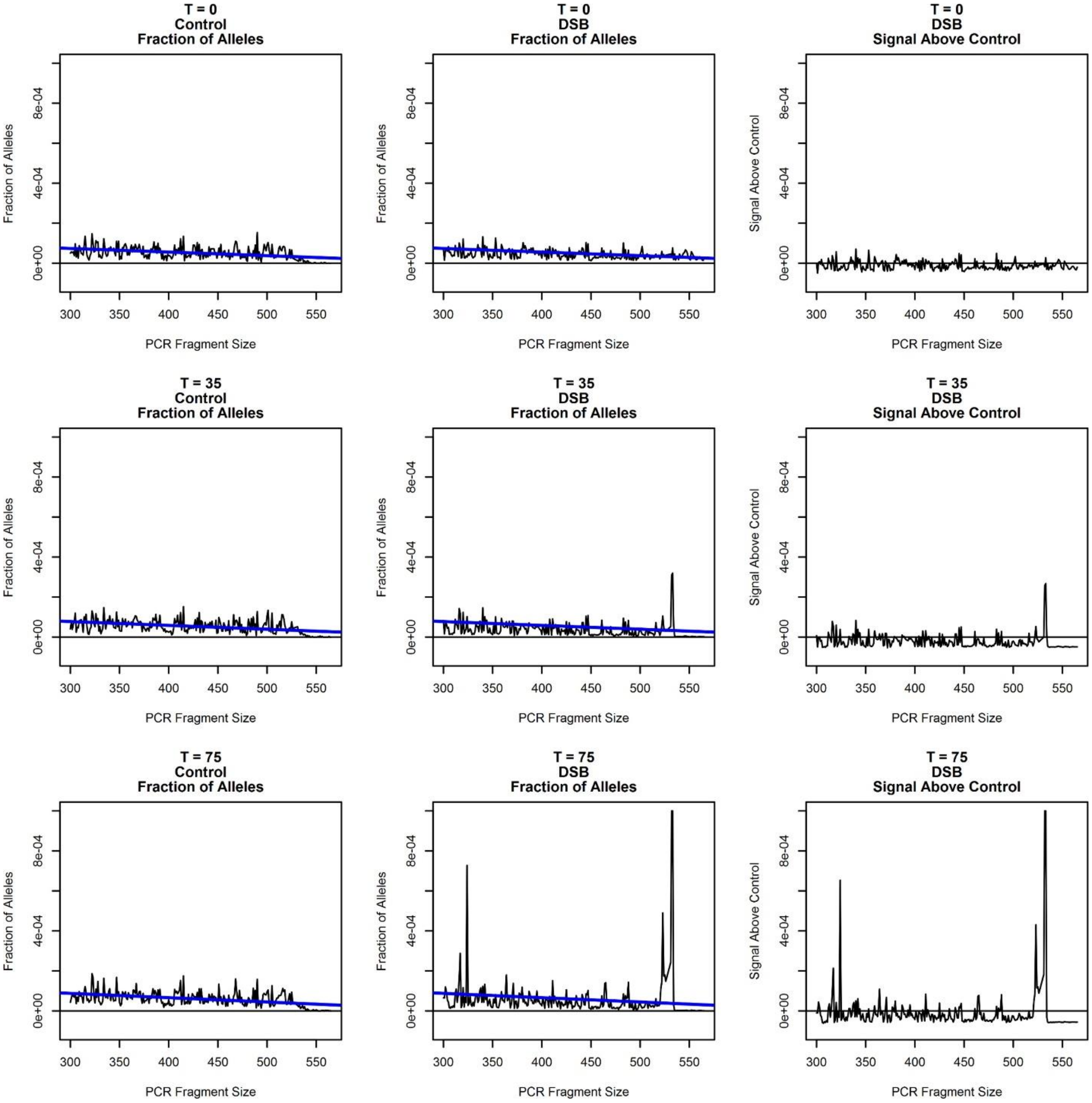


**Supplementary Figure 4: Demonstration of data normalization.**
Nine plots show three data traces for each of three exemplary time points (T0, before DSB induction; T35, just after DSB formation; T75, in active resection). For each time point, the first two plots show the calculated fraction of all input alleles giving sequencing reads of the indicated PCR fragment sizes (i.e. allele positions), excluding the DSB and NdeI end positions. The blue line is a linear regression fit to the control data; that same line is drawn on the control (left) and DSB (middle) plots. Note the small negative slope that reflects PCR amplification bias. The rightmost plot in each row shows the DSB allele fraction minus the linear regression fit, referred to as “signal above control”. Note that at T0, the signal above control averages to zero and that peaks only appear after resection ensues, validating the approach.

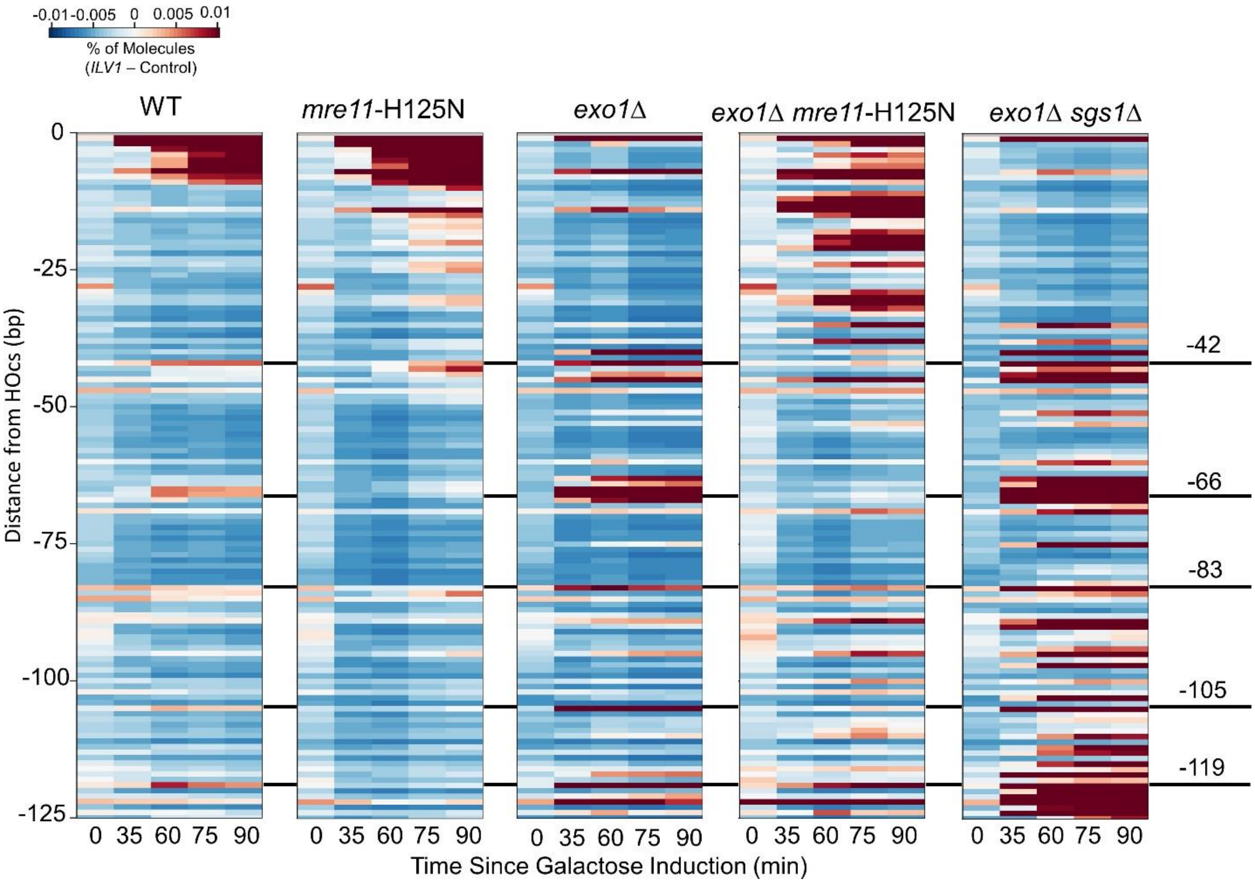


**Supplementary Figure 5: Evidence for multiple incision positions with variable efficiency.** Five heatmaps depict the number of unique *ILV1*-PR molecules at positions -1 to -125 bp from the HOcs in WT and various mutants. Positions with increases in unique reads dependent on Mre11 and time after DSB induction are aligned and labeled to facilitate comparison. The higher plotting sensitivity is the same as used in Figures 1G and 3A. Note that heat map coloring can be misleading when saturated; comparison to scatterplots is recommended.

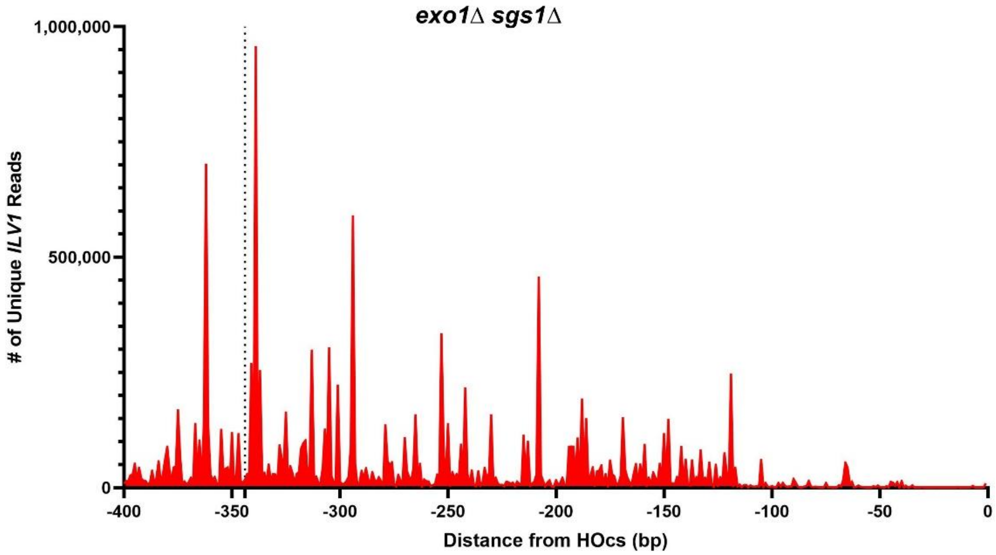


**Supplementary Figure 6: Resection in *exo1*Δ *sgs1*Δ mutant yeast continues past -350 bp.** It was possible to examine resection into the common sequence that was shared between the *ILV1* and control alleles for the *exo1*Δ *sgs1*Δ strain because the signal was dramatically higher that the background fragmentation of the control allele. However, our typical normalization approach could no longer be used. For this plot we simply aligned reads to the DSB allele and summed the total read counts that aligned for the T60, T75 and T90 minute time points from two independent *exo1*Δ *sgs1*Δ experiments. The dashed line at position -344 bp marks the boundary between sequences unique to the DSB allele and those shared with the control allele. The large resection signal continues past this point throughout the possible range of PCR products.

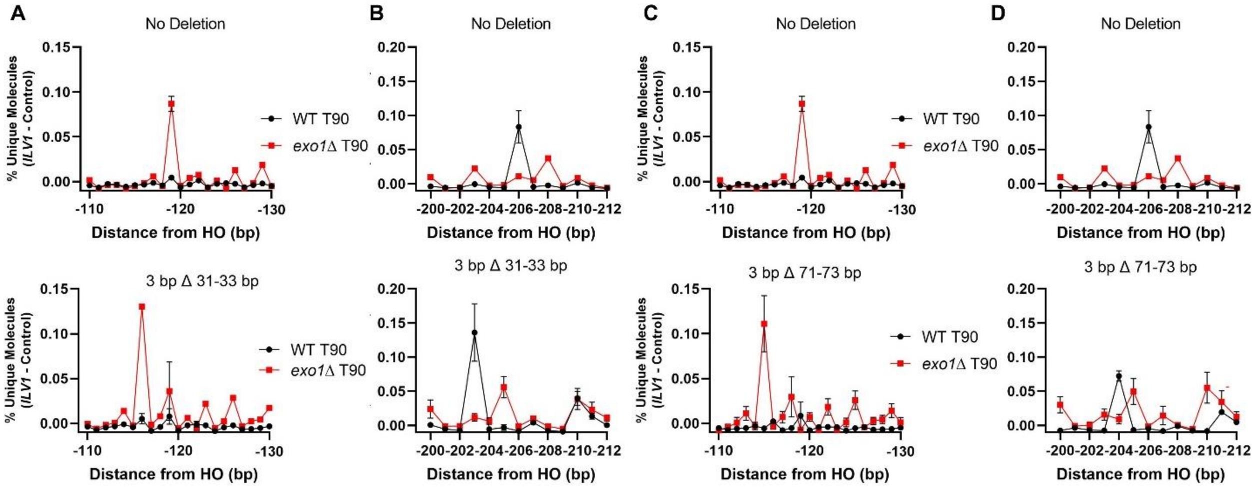


**Supplementary Figure 7: Three base deletions shift the entire distal resection pattern.** **(A&C)** The signal around the -119 bp peak was plotted in WT and *exo1*Δ strains with and without the 3 bp deletion either **(A)** 31-33 bp or **(C)** 71-73 bp from the HOcs. **(B&D)** The same as (A&C) for the region surrounding the -206 bp peak. In all cases, the peak patterns moved 3 bp closer to the DSB, demonstrating their sequence specificity.

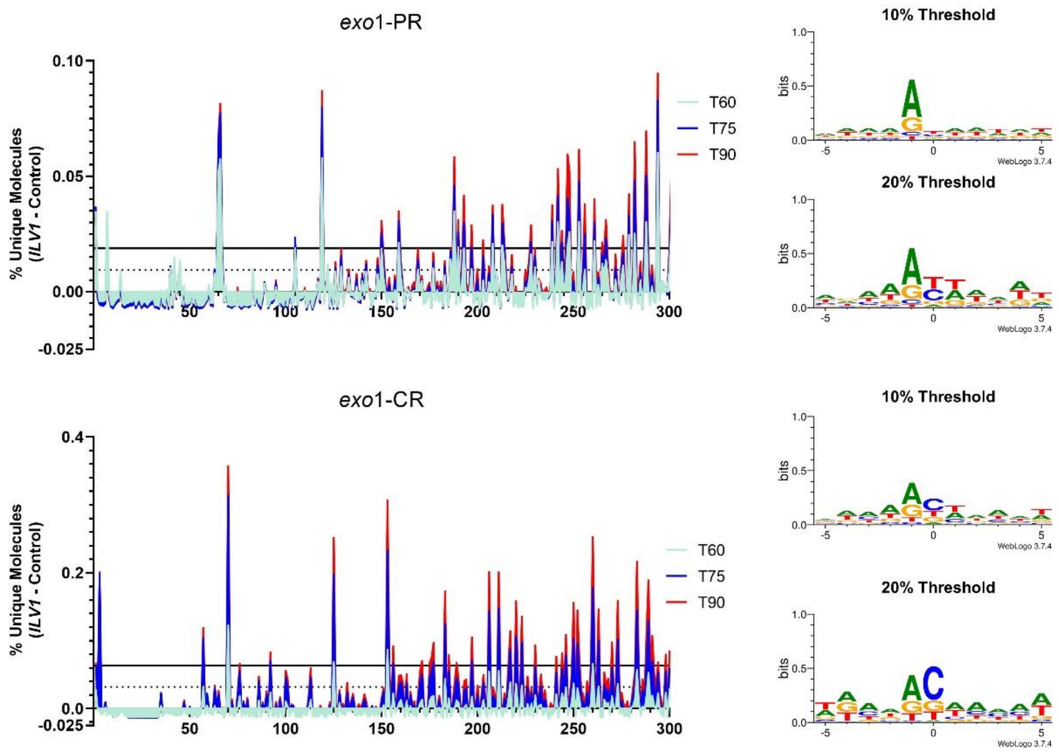


**Supplementary Figure 8: Depiction of logo plot thresholds.**
Shaded line graphs show resection signal in *exo1* and *exo1 sgs1* mutant yeast similar to Figure 4D with horizontal lines denoting the thresholds used for generating logo plots. Dotted lines are at 10% of max peak height as used to make logo plots in Figure 6E (repeated in this figure) while solid lines are at 20% of max peak height as used to make logo plots in this figure.

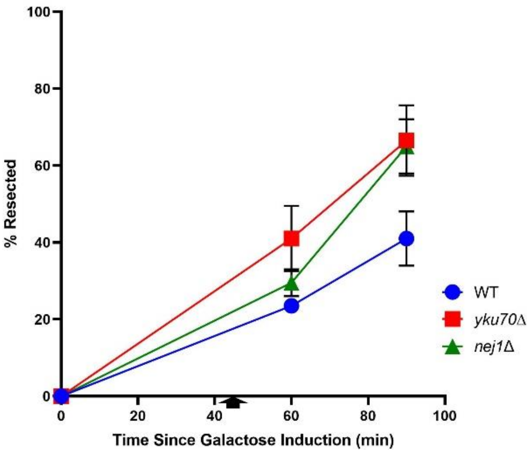


**Supplementary Figure 9: Deletion of *yku70* or *nej1* increases the resection rate.**
ddPCR-based resection assay to determine the percentage of ssDNA after DSB induction at a site 355 bp away from the HOcs in WT (also shown in Figure 1B), *yku70*Δ and *nej1*Δ yeast strains. Cells were washed into glucose at 45 minutes (arrow).

**-65 position (prominent in WT, Mre11-dependent)**

```
3’ - TCGAAAGGCGTTGTCATATTAAAA - 5’ HOcs, orientation #1
     |         ||  | | ||
3’ - TTTTTTATTCTTCCCGTTTTTCTT - 5’ possible HOcs homology?
     ||||    |     ||  ||||
3’ - TTTTAATATGACAACGCCTTTCGA - 5’ HOcs, orientation #2
     ---------*
```

**-66 position (prominent in WT, Mre11-dependent)**

```
3’ - TCGAAAGGCGTTGTCATATTAAAA - 5’ HOcs, orientation #1
                |  |   ||
3’ - CTTTTTTATTCTTCCCGTTTTTCT - 5’ possible HOcs homology?
      |||  |||     |   |||
3’ - TTTTAATATGACAACGCCTTTCGA - 5’ HOcs, orientation #2
     ---------*
```

**-206 position (prominent in WT, Exo1-dependent)**

```
3’ - TCGAAAGGCGTTGTCATATTAAAA - 5’ HOcs, orientation #1
       ||  | ||  |   | | |  |
3’ - GTGATCGCCGAGGAACTTTAAGTA - 5’ possible HOcs homology?
      |       ||  |    |    |
3’ - TTTTAATATGACAACGCCTTTCGA - 5’ HOcs, orientation #2
     ---------*
```

**-294 position (prominent in *exo1 sgs1*)**

```
3’ - TCGAAAGGCGTTGTCATATTAAAA - 5’ HOcs, orientation #1
        |||||   |     |    ||
3’ - AACAAAGGGCGTTGTCGACGTTAA - 5’ possible HOcs homology?
         ||              |  |
3’ - TTTTAATATGACAACGCCTTTCGA - 5’ HOcs, orientation #2
     ---------*
```

**-294 position (prominent in *exo1 sgs1*) ALTERNATIVE ALIGNMENT REGISTER**

```
3’ -  TCGAAAGGCGTTGTCATATTAAAA - 5’ HOcs, orientation #1
       | || |||||||||    | ||
3’ - AACAAAGGGCGTTGTCGACGTTAA  - 5’ possible HOcs homology?
          |    |    || | ||
3’ -  TTTTAATATGACAACGCCTTTCGA - 5’ HOcs, orientation #2
     ---------*
```

**Supplementary Figure 10: The -294 bp position has sequence similarity to the HOcs.**
To query whether HO might be cleaving the *ILV1*-PR sequence in an off-target fashion to lead to some signal peaks, we aligned the sequence surrounding key resection intermediates (middle rows) to both orientations of the HOcs including the putative 3’ overhangs left by HO (green) and what would be mapped as the 5’ terminal position if HO had cut (red), as compared to the actual mapped 5’ position (blue). None were convincing as possible imperfect HOcs sequences as aligned, but we noted that the -294 peak observed with *exo1 sgs1* yeast had a possible HOcs alignment if we shifted it 1 bp from the actual cleavage position.

### Supplementary Tables

### Supplementary Table 1: Yeast strains used in this study.

| Strain # | Genotype |
|---|---|
| YW3104 | *MATa-inc::LEU2 bar1Δ::KanMX4 dnl4::K466A gal1::HO his3Δ1 ILV1::ori-his3MX6 ILV1prm::HOcs leu2Δ0 met15Δ0 ura3Δ0* |
| YW3177 | YW3104 + ChrXIII::ILV1CtrlAllele-Ura3 |
| YW3185 | YW3104 + *exo1Δ::HYG* |
| YW3189 | YW3104 + *yku70ΔMET15* |
| YW3193 | YW3104 + *sgs1Δ::HYG* |
| YW3197 | YW3104 + *sae2Δ::HYG* |
| YW3201 | YW3193 + ChrXIII::ILV1CtrlAllele-Ura3 |
| YW3203 | YW3185 + ChrXIII::ILV1CtrlAllele-Ura3 |
| YW3205 | YW3197 + ChrXIII::ILV1CtrlAllele-Ura3 |
| YW3207 | YW3189 + yku70ΔMET15 |
| YW3213 | YW3104 + *tel1Δ::HYG* |
| YW3219 | YW3104 + *mre11-H125N* |
| YW3222 | YW3219 + *mre11-H125N* |
| YW3229 | YW3104 + *ILV1::URA3*ins31bp |
| YW3231 | YW3104 + *nej1ΔHYG* |
| YW3232 | YW3231 + ChrXIII::ILV1CtrlAllele-Ura3 |
| YW3236 | YW3104 + *mre11*Δ::*HYG* |
| YW3237 | YW3236 + ChrXIII::ILV1CtrlAllele-Ura3 |
| YW3246 | YW3104 + *ILV1::URA3*ins71bp |
| YW3250 | YW3213 + ChrXIII::ILV1CtrlAllele-Ura3 |
| YW3254 | YW3229 + *ILV1::*3bpΔ31-33 |
| YW3256 | YW3246 + *ILV1::*3bpΔ71-73 |
| YW3262 | YW3185 + *mre11-H125N* |
| YW3263 | YW3262 + ChrXIII::ILV1CtrlAllele-Ura3 |
| YW3266 | YW3254 + ChrXIII::ILV1CtrlAllele-Ura3 |
| YW3268 | YW3256 + ChrXIII::ILV1CtrlAllele-Ura3 |
| YW3274 | YW3254 + *exo1Δ::HYG* |
| YW3275 | YW3256 + *exo1Δ::HYG* |
| YW3276 | YW3274 + ChrXIII::ILV1CtrlAllele-Ura3 |
| YW3277 | YW3275 + ChrXIII::ILV1CtrlAllele-Ura3 |
| YW3286 | YW3185 + *sgs1*Δ::*NAT* |
| YW3287 | YW3286 + ChrXIII::ILV1CtrlAllele-Ura3 |
| YW3289 | YW3104 + ChrXV::ILV1-CR_CtrlAllele-URA3 |
| YW3291 | YW3185 + ChrXV::ILV1-CR_CtrlAllele-URA3 |

### Supplementary Table 2: Oligonucleotides used in this study.

| Oligo # | Type | Modifications | Sequence (5' to 3') | Purpose |
|---|---|---|---|---|
| OW3908 | Primer | | TCCACCACGATACGGGAAACAG | ddPCR *ILV1*-PR Spike-In % |
| OW3991 | Primer | | AATAAGAAGGGCAAAAAGAAAAAGC | HOcs ddPCR |
| OW3992 | Primer | | AAAGCAGCAACAACAAAAGTTTTC | HOcs ddPCR |
| OW3995 | Primer | | AACCTTCCACATATCGTCCAATTC | ddPCR Resection Monitoring |
| OW4079 | Primer | | TGCGGGAAACAAATATCTTGAGG | ddPCR Resection Monitoring |
| OW4090 | Probe | FAM | ATGGGTCCTTGGATTCTCGCTAAACAGTCT | ddPCR Resection Monitoring |
| OW4161 | Primer | | GCCGGTATTGACCAAACTACTT | *ACT1* Control |
| OW4162 | Probe | VIC | TTATACGGTAACATCGTTATGTCCGGTGGT | *ACT1* Control |
| OW4163 | Primer | | TCATGGAAGATGGAGCCAAA | *ACT1* Control |
| OW4221 | Probe | FAM | CGCTTTTAGTTTCAGCTTTCCGCA | HOcs ddPCR |
| OW4282 | Primer | 12 bp UMI | TAGACTGATNNNNNNNNNNNNTGATCCAAAAAGCCAACACCTGTGT | *ILV1*-PR Primer Extension |
| OW4283 | Primer | 12 bp UMI | TACTAGTCTNNNNNNNNNNNNTGATCCAAAAAGCCAACACCTGTGT | *ILV1*-PR Primer Extension |
| OW4284 | Primer | 12 bp UMI | TATGCATGTNNNNNNNNNNNNTGATCCAAAAAGCCAACACCTGTGT | *ILV1*-PR Primer Extension |
| OW4285 | Primer | 12 bp UMI | TAGCTACATNNNNNNNNNNNNTGATCCAAAAAGCCAACACCTGTGT | *ILV1*-PR Primer Extension |
| OW4286 | Primer | 12 bp UMI | TACACGTATNNNNNNNNNNNNTGATCCAAAAAGCCAACACCTGTGT | *ILV1*-PR Primer Extension |
| OW4287 | Primer | 12 bp UMI | TATCAGACTNNNNNNNNNNNNTGATCCAAAAAGCCAACACCTGTGT | *ILV1*-PR Primer Extension |
| OW4288 | Primer | | TCGTCGGCAGCGTCAGATGTGTATAAGAGACAGTTCCCAAGCTACTCAGTCACAG | *ILV1-PR* Library Amplification |
| OW4290 | Primer | | GTCTCGTGGGCTCGGAGATGTGTATAAGAGACAGAACCTTCCACATATCGTCCAATTC | *ILV1-PR* Library Amplification |
| OW4294 | Probe | FAM | CTAAACCCAGTTCAACATCACCACA | ddPCR *ILV1*-PR Spike-In % |
| OW4299 | Primer | | GAATAAGAAGGCAGAAAGGGTAAGG | ddPCR *ILV1*-PR Spike-In % |
| OW4382 | Primer | | ACACCTTGAGAGATTGGAGATT | ddPCR *ILV1*-CR Spike-In % |
| OW4383 | Primer | | ATCGAGACCAATAAGAACATTGC | ddPCR *ILV1*-CR Spike-In % |
| OW4384 | Probe | FAM | TCAGTTTGCAGCTCATCCATGT | ddPCR *ILV1*-CR Spike-In % |
| OW4388 | Primer | | TCGTCGGCAGCGTCAGATGTGTATAAGAGACAGATGGTACGTCTGTGCGTATG | *ILV1*-CR Library Amplification |
| OW4389 | Primer | | GTCTCGTGGGCTCGGAGATGTGTATAAGAGACAGACACCTTGAGAGATTGGAGATT | *ILV1*-CR Library Amplification |
| OW4401 | Primer | 10 bp UMI | TAGACTGATNNNNNNNNNNNNAACAAGAACAACTTCATCAACCACT | *ILV1*-CR Primer Extension |
| OW4402 | Primer | 10 bp UMI | TACTAGTCTNNNNNNNNNNNNAACAAGAACAACTTCATCAACCACT | *ILV1*-CR Primer Extension |
| OW4403 | Primer | 10 bp UMI | TATGCATGTNNNNNNNNNNNNAACAAGAACAACTTCATCAACCACT | *ILV1*-CR Primer Extension |
| OW4404 | Primer | 10 bp UMI | TAGCTACATNNNNNNNNNNNNAACAAGAACAACTTCATCAACCACT | *ILV1*-CR Primer Extension |
| OW4405 | Primer | 10 bp UMI | TACACGTATNNNNNNNNNNNNAACAAGAACAACTTCATCAACCACT | *ILV1*-CR Primer Extension |
| OW4406 | Primer | 10 bp UMI | TATCAGACTNNNNNNNNNNNNAACAAGAACAACTTCATCAACCACT | *ILV1*-CR Primer Extension |
